## Supporting Information for "Whole-chromosome hitchhiking driven by a male-killing endosymbiont"

### S1 Text: Supplementary Methods

#### Reference genome sequencing and assembly

An adult female from the Nairobi area (RF.K001) was used to generate short insert libraries (500 bp) for paired end sequencing, and another female from the same brood was used to generate mate-pair libraries (insert sizes of 2 kb, 4.2 kb, 8.5 kb, 9.5 kb and 11.5 kb). Sequencing was performed using Illumina HiSeq 2500 technology with a read length of 250 bp. In total more than half a billion reads (336,807,664 x 2 = 673,615,328) containing more than 892 Gb (892,999,615,400 bp) of raw sequence generated from these two sisters (S1 Table).

Read files were quality checked using FastQC v0.10.1 [1]. Reads from small insert libraries were pre-processed with Trimmomatic v0.33 [2] to remove external adapter contamination as well as a minimum length threshold of 100 bp to discard false mate pair reads (ILLUMINACLIP::2:7:7 MINLEN:100).

All adapter-trimmed reads were checked for possible contamination using FastqScreen v0.5.2 [3] with libraries from human (*Homo sapiens* GRCh38), mouse (*Mus musculus* GRCm38), *E. coli* (U00096.3), Enterobacteria phage phiX174 (NC_001422.1), *Danaus chrysippus* mt genome as positive control (NC_026538.1) and simulated bacterial and viral databases from DeconSeq [4]. Most sequences could not be mapped to the provided libraries. Small fractions of reads could be mapped multiple times to multiple genomes which hint towards similar repeats in the provided libraries.

Genome size and overall genome characteristics (total and haploid genome length, percentage of repetitive content, and heterozygosity) were estimated from the k-mer profile using GenomeScope v1.0.0 [5]. This indicated high genome heterozygosity with a bi-modal k-mer distribution where the frequency of heterozygous k-mers (peak at 75x) were higher than homozygous k-mers (peak at ~150x). Inferred genome characteristics are given in S2 Table.

The k-mer profile of *Danaus chrysippus* suggested a high level of heterozygosity of about 3%. We therefore adopted a multi-step genome assembly pipeline designed to account for this heterozygosity. An initial draft assembly was generated by a single run of the SPAdes v3.8.1 [6] assembler using default parameters. For de-novo assembly, SPAdes selects multiple k-mer sizes, such as 21, 33, 55, 77, 99 and 127. Additional mate-pair data were incorporated to further scaffold the diploid assembly. We then used Redundans v0.12a [7] to generate a haplotype resolved and scaffolded assembly. Scaffolding was performed using default parameters with all six (PE + MP) libraries in ascending order of average insert size according to the estimated ELF fraction. All sequences smaller than 500 bp were excluded from subsequent assembly steps. We iterated the Redundans runs multiple times to utilize reads with different orientations as a result of PE contamination in Nextera mate-pair data. Using the iterative approach we managed to increase the N50 of the assembly from 20kb to 204kb. However, the assembly size was still much higher than the 250 Mb (non-repetative fraction) estimated from the k-mer profile. We therefore used Haplomerger2 v3.4.0 [8,9], to further refine the scaffolds using mate-pair data with default parameters. Finally, a single scaffold of less than 1 kb was discarded and the mitochodrial scaffold and one contaminant scaffold were removed, to produce our final assembly. The final assembly N50 is 628 kb and the total assembly size is 322 Mb. Detailed characteristics are provided in S3 Table.

**Repeat content analysis**

A repeat library was created using dnaPipeTE 1.2 [10] and RepeatModeler 1.0.4 (http://www.repeatmasker.org/RepeatModeler/). For dnaPipeTE, mitochondrial and forward reads were excluded (as per the manual) from the complete trimmed raw read dataset, thereby producing an input file containing only reverse and unpaired reads from all libraries (168733627 reads / 42,183,406,750 bp). Firstly, dnaPipeTE was run on 30 coverage points ranging from 0.0006 to 0.8 with an estimated genome size of 400 MB. All parameters were set as default except the minimal contig length which was lowered from 200 to 50 bp. After analysing the N50 distribution from 30 coverage points, a coverage of 0.03 was selected as an optimal sampling coverage. 50 repetitions were performed with the selected optimal coverage and the same parameters as above. This resulted in 99,564 contigs with total length of 41,355,997 bp. The maximized N50 at 0.009 coverage was not viewed to be the optimal sampling coverage because the samples are very small and a huge variation in N50 was expected. Furthermore, filtering steps were introduced afterwards to discard non-repeats.

All trimmed reads were mapped un-paired against all contigs from dnaPipeTE using BWA mem v0.7.12 [11,12] with the options -t 80 -k 25 -a -y 26 -c 1000000000 and otherwise default settings. The coverage per position was calculated using samtools v1.3 [13] mpileup with options -A -C 50 -d 1000000 and otherwise default parameters. Contigs with a median coverage smaller than the 90% quantile (94x) of the per position coverage distribution were filtered out (3,876 out of 99,564 contigs). The remaining 95,688 (40969029 bp total length) with sufficient coverage were used in the subsequent steps.

A repeat-protein free protein database was built from Swiss-Prot database [14] (accessed on March 30th 2017) and all repeat sequences from the Repbase database [15] (accessed on March 30th 2017). The repeat sequences were searched in the Swiss-Prot database via BlastX 2.3.0 [16,17] with an e-value cutoff of e10-11. Protein sequences with hits from repeat sequences containing at least 20 bp in one hit were removed to obtain a repeat-protein free protein database. The repeat families from dnaPipeTE runs were blasted (BlastX; e-value cutoff e10-11) against the repeat-protein free protein database. These sequences were removed from final repeat library containing 41261 sequences totalling a length of 17477008 and an N50 of 736 bp.

RepeatModeler was then executed with default parameters on the final genome assembly and on the contigs obtained from dnaPipeTE. The resulting fasta files containing the repeat families were concatenated into 936 sequences with a total length of 1,433,744 bp and an N50 of 2644 bp.

**Assembly Completeness**

The Core Eukaryotic Gene Mapping Algorithm, CEGMA v2.5.0 [18] and the Benchmarking Universal Single-Copy Orthologs, BUSCO v3.0.0 [19] were used to estimate the completeness of genome assembly and quality of gene annotation of the *D. chrysippus* final assembly. Results are presented in S4 Table and S5 Table. We ran BUSCO with bot the Arthropoda (n=1066) and Insecta (n=1058) core gene sets which identified 93% and 94% of the single copy genes searched, respectively. This suggest a high contiguity and completeness of our final assembly.

**Genome Annotation**

For annotation of the assembly we used the MAKER v2.31.8 pipeline [20,21] combined with MPICH2 (http://www.mpich.org/) in three iterations. Firstly, an Augustus species model was computed using the gene models obtained from BUSCO analysis. BUSCO was run on the assembly using the metazoan dataset together with the option --long and otherwise default parameters. From the CEGMA run on the assembly a SNAP [22] model was built using the script cegma2zff from the MAKER2 distribution and the SNAP scripts fathom (fathom genome.ann genome.dna -categorize 1000 && fathom -export 1000 -plus uni.ann uni.dna), forge (export.ann export.dna) and hmm-assembler.pl. A Genemark [23] model was built using GeneMark-ES suite 4.32 from self-training (--ES) on the assembly. An initial MAKER2 run was executed using the *D. chrysippus* assembly, *D. chrysippus* ESTs (obtained from GenBank), and protein sequences from *Danaus plexippus* (as a protein homology evidence from an alternate closely related species) with options est2genome and protein2genome enabled. In the second MAKER2 run, the est2genome and protein2genome options were disabled and the *D. chrysippus* assembly was used as input, along with the Augustus species model, custom repeat library HMM models from SNAP and GeneMark. The minimum protein length was set to ten amino acids. After the first iteration, the gff file for the whole assembly was extracted using the MAKER2 gff3_merge, converted with maker2zff and a new HMM model was built for SNAP the same way as above. The Augustus species model was retrained locally by first converting with the SNAP script zff2gff3.pl (zff2gff3.pl genome.ann | perl -plne 's/\t(\S+)$/\t\.\t$1/') and second the autoAug.pl script from Augustus 3.2.2 [24]. The input for the autoAug.pl were the genome assembly, trained Augustus species model and the gff3 file created from the first MAKER2 iteration. Default parameters wre used, but with the options -v --useexisting. For the second MAKER2 iteration the SNAP HMM model was further improved using the results from first iteration. Additionally, the Augustus species model was also improved. At this point, the minimum protein length was raised to 30 amino acids. Thereafter, a retraining as above and a third MAKER2 run was performed. Functional annotation of the MAKER2 derived gene models were done using homology searches against BLAST nr, InterPro and KEGG databases. A summary of the annotation characteristics is provided in S6 Table.

**Orthology analysis**

Protein sets from *Danaus chrysippus*, three other butterfly species and a moth species were used to predict ortholog clusters using OrthoFinder v0.7.1 [25] with default parameters. Functional annotation of all protein sets with Gene Ontology (GO) terms was performed with BLAST2GO v4.1.0 [26] and InterProScan v5.0.0 [27] using default parameters. Results are presented in S7 Table S7 and S8 Table.

**Pseudo-chromosomal assembly**

We produced a pseudo-chromosomal assembly based on homology with the *Heliconius melpomene* Hmel2.5 genome, which is the most contiguous genome from the same family (Nymphalidae) currently available. Following [28], we numbered chromosomes based on the M.

cinxia genome [29]. *Danaus* have 30 chromosomes, which differ from the ancestral 31 (as found in *M. cinxia*) by a single fusion between chromosome 1 (the Z sex chromosome) and chromosome 21 [28]. However, *Heliconius melpomene* has 21 chromosomes due to 10 fusion events [29,30]. In order to assign *Danaus* chromosomes for each gene, we therefore first determined the tracts representing the ancestral 31 chromosomes in the ancestor of *Heliconius* by splitting at the fusion points identified by Davey et al. [30], re-numbering the chromosomes based on *M.* cinxia [29], and then fusing chromosomes 1 and 21 to effectively reorganise the *Heliconius* chromosomes into a *Danaus*-like karyotype..

We used BLASTp [16,17] to identify likely homology between the translated gene sets for the two species. Only hits with an e value < $10^{-20}$ and a minimum AA sequence identity of 50% were considered. We first screened for misassembled scaffolds by identifying all scaffolds with hits to two or more chromosomes, with the additional requirement that at least three genes (which had to represent at least 5% of genes on the scaffold) had hits to the lesser chromosome. All possible misassembled scaffolds were then inspected visually and likely breakpoints were identified in 80 scaffolds. The resulting modified assembly consisting of 945 scaffolds was used for all evolutionary analyses.

To produce a pseudo-chromosomal assembly, we discarded all scaffolds with fewer than four blast hits to a single chromosome except for those in which at least 75% of blast hits were to a single chromosome. In total 438 scaffolds, totalling 282 Mb (87% of the genome) could be assigned to chromosomes. Scaffolds were ordered based on the median position of their blast hits, and oriented based on the rank order of the start and end positions of all hits.

##### *Spiroplasma* genome

Scaffolds corresponding to the genome of the *Spiroplasma* sp. endosymbiont were identified based on comparative read depth in suspected infected and uninfected samples (S10 Fig). Initially, only a single *Spiroplasma* scaffold was identified in the complete genome, but inspection of the second version of the genome (i.e. before running Haplomerger, see above) identified twelve scaffolds with high read depth in the infected sample and effectively zero read depth in the cured sample. Three scaffolds appeared to represent one or more *Spiroplasma* plasmids, as their read depth was considerably higher. One of the other nine scaffolds was evidently chimeric based on read depth, and this scaffold was broken manually. The resulting putative *Spiroplasma* genome (accession XXX) has a total length of 1.7 Mb. We annotated the genome using RAST [31,32], and confirmed its identity using BLAST [17] to available *Spiroplasma* genomes. All but the shortest scaffold had multiple hits to *Spiroplasma sp.* (e vlaue < 1e-10). The other two candidate plasmid scaffolds had hits to *Spiroplasma citri* plasmids. To test for infection in resequenced samples, reads were mapped to our *Spiroplasma* sp. genome using bwa mem v0.7.12 [11,12] and read depth was computed using Samtools v1.3 [13]. We also confirmed that other known male-killers *Wolbachia* and *Rickettsia* are not represented in our reference genome (from a member of an all female brood) using blast with published sequences (GenBank accessions AJ130716 and AJ269519).

**Mitochondrial genome assemblies**

We assembled the mitochondrial genomes for all 45 samples included in the study from Illumina short-read sequences using NOVOplasty [33]. A previously sequenced *D. chrysippus* mitochondrial genome (GenBank accession: NC_024532) was used as a seed. Although NOVOplasty generates de novo assemblies, a reference sequence can be provided to resolve duplicated regions. The same complete genome was therefore also provided as a reference. For all 42 *D. chrysippus* samples sequenced in this study, which were either 150 bp paired-end or 250 bp paired-end, we used a kmer size of 39, an insert size of 300 and an insert range of 1.8 (1.3 for repetitive regions). For the *D. petilia* and *D. gilippus* samples that were sequenced previously [34] (100 bp paired-end), we used a kmer size of 25, an insert size of 300 and an insert range of 2.4 (1.3 for repetitive regions).

**Population sample resequencing and genotyping**

DNA was extracted from thorax tissue using the Dneasy blood and tissue kit (Qiagen, ). Paired end sequencing libraries were prepared using the Truseq Nano DNA HT Sample preparation Kit (Illumina USA) with the addition of individual indexes. Fragmentation to 350 bp was performed using the Covaris cracker. Fragments were prepared for Illumina sequencing via adapter ligation and further PCR amplification, followed by purification using the AMPure XP system (Beckman Coulter). Paired-end sequencing (150 bp) was performed using the HiSeq X (Illumina). All samples were sequenced to a mean depth of coverage 20x or greater.

Reads were mapped to the *D. chrysippus* reference assembly using Stampy [35] v1.0.31. BAM files were sorted using Samtools [13] v1.3 and PCR duplicate reads were removed using PicardTools MarkDuplicates v1.135 (https://broadinstitute.github.io/picard/). Genotyping was performed using GATK v3 HaplotypeCaller and GenotypeGVCFs [36,37] using default parameters except that heterozygosity was set to 0.02. Genotyping was performed separately for each species. Genotype calls were required to have an individual depth ≥ 8x, and heterozygous and alternate allele calls were further required to have an individual genotype quality (GQ) ≥ 20.

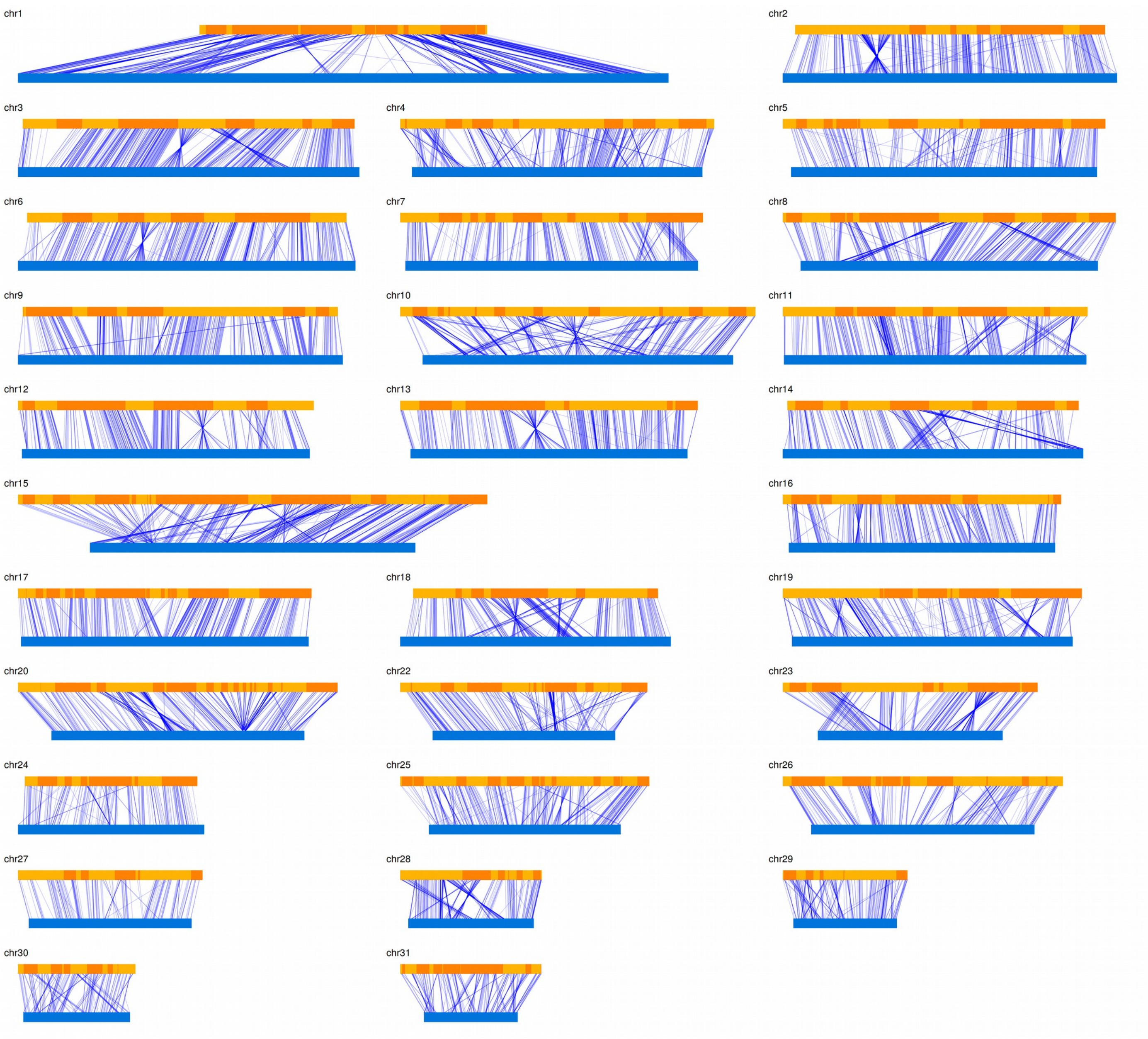


**S1 Fig. Pseudo-chromosomal assembly of *D. chrysippus.*** Homology with the *Heliconius melpomene* genome (corrected for known fusion events [28–30] and scaffolded into chromosomes [38]) (blue) allowed us to construct a robust pseudo-chromosomal assembly for *D. chrysippus*. Scaffolds of *D. chrysippus* are shown in alternating shades of orange. Blue lines connect homologous genes (Blast e-value < 1e-20, identity > 50%).

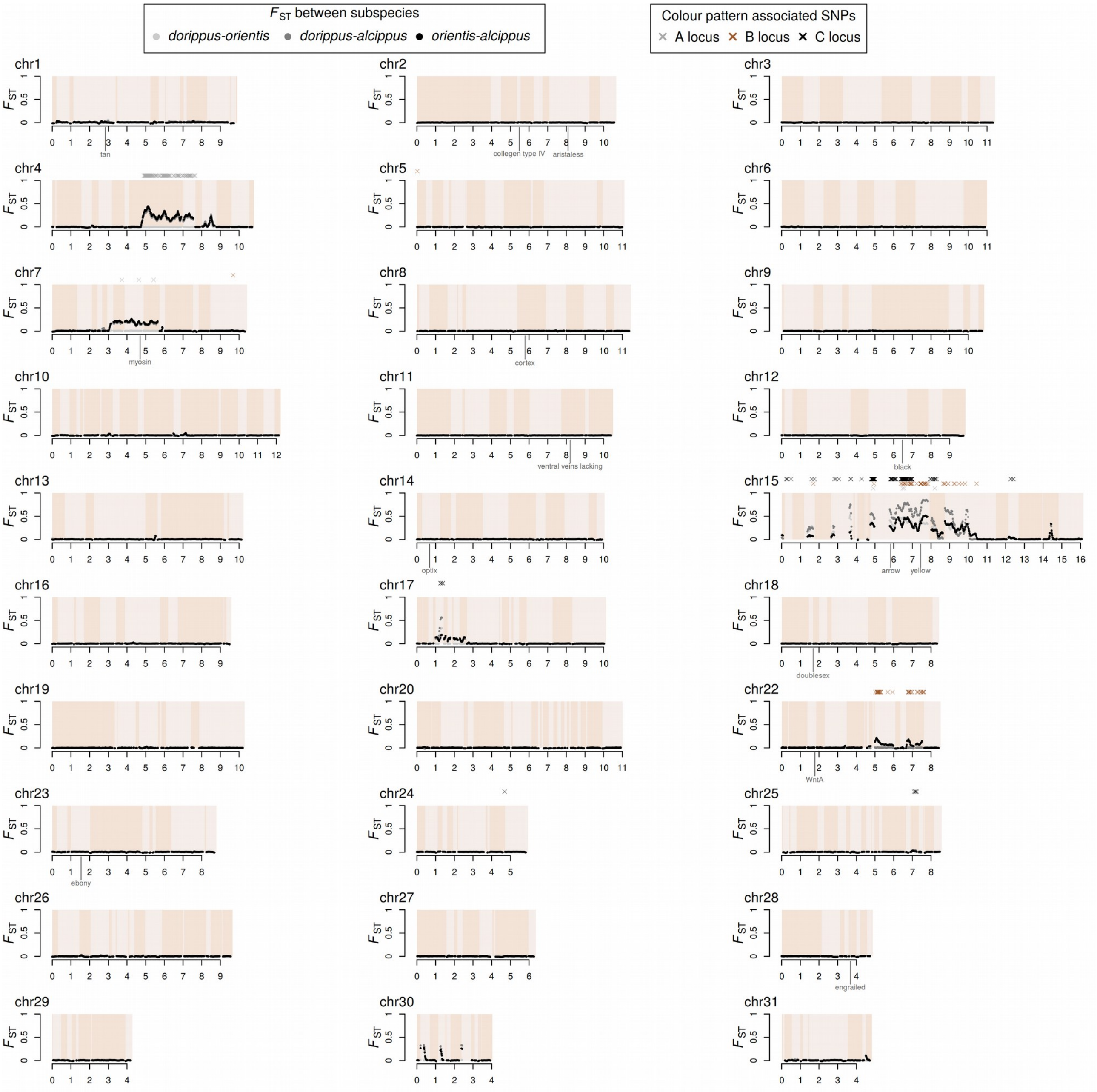


**S2 Fig. Genetic differentiation and SNP associations with colour pattern.** $F_{\mathrm{ST}}$ is plotted across each chromosome between three different subspecies of *D. chrysippus*, as indicated above the plot. Scaffolds are indicated by light and dark shading. Numbers on the x-axis indicate chromosome position in Mb. Coloured crosses above the plots indicate SNPs strongly associated with the phenotypes controlled by the A (blue), B (brown) and C (black) loci (Wald test, 99.99% quantile). A number of candidate genes are annotated on the plot. These include known and putative wing patterning genes in *Heliconius* (*optix* [39], *cortex* [40], *WntA* [41]*, aristaless* [42] *and ventral veins lacking* [43]) and *Papilio* spp. (*doublesex* [44] and *engrailed* [45]). A *myosin* gene thought to be associated with a pale mutant form in *Danaus plexippus* [34] is also indicated, along with *collegen type IV*, which was found to be associated with migratory behaviour in *D. plexippus* [34]. Several melanism-related genes are also annotated, as well as *arrow* which was added to the list of candidates post-hoc due to strong association with colour pattern (see main text). Of our *a priori* candidates, only *yellow* is found to associate with colour pattern in *D. chrysippus*.

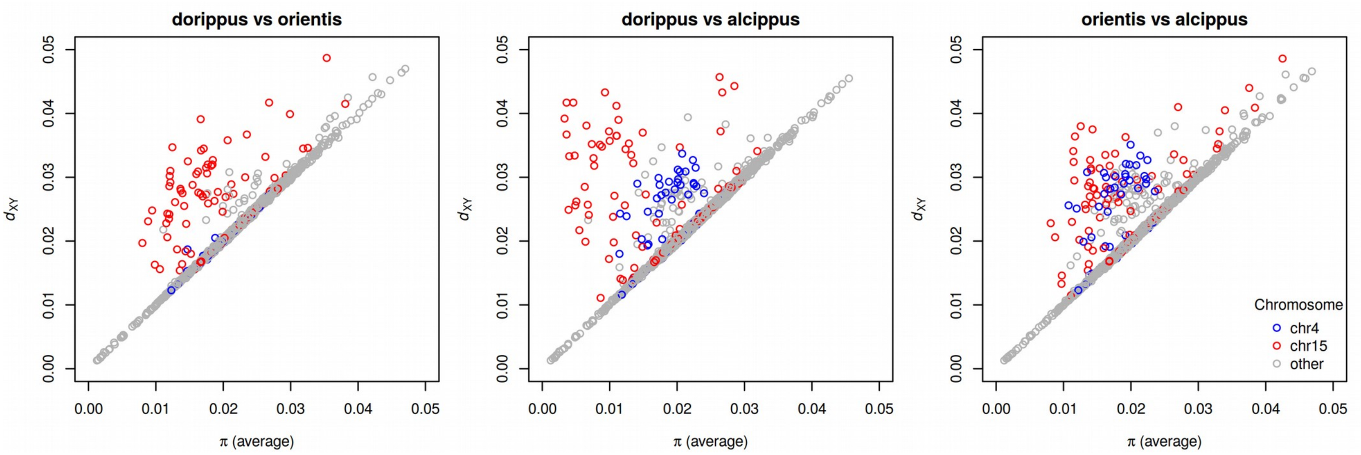


**S3 Fig. $d_{XY}$ plotted against π reveals excess divergence on colour pattern-associated chromosomes.** Absolute divergence between each pair of populations ($d_{XY}$) and nucleotide diversity within populations (π) were computed for non-overlapping 100 kb windows. The value of π plotted is the average between the two populations in each plot. The clustering of points along the diagonal indicate that diversity within each subspecies is similar to divergence between subspecies, consistent with a single nearly-panmictic population. Points that deviate to the left of the diagonal indicate either excess divergence between subspecies or reduced diversity within subspecies, or both. Here, the colour-pattern associated regions on chromosome 4 and 15 (indicated in colour for convenience) show signatures of local adaptation with both reduced within-population diversity and increased between-population divergence, as would be expected if selection limits effective gene flow at these loci. One pair of populations, *dorippus* and *orientis*, are diverged at chromosome 15 but not chromosome 4, which is also expected as they only differ in their forewing phenotype and both lack the white hindwing patch.

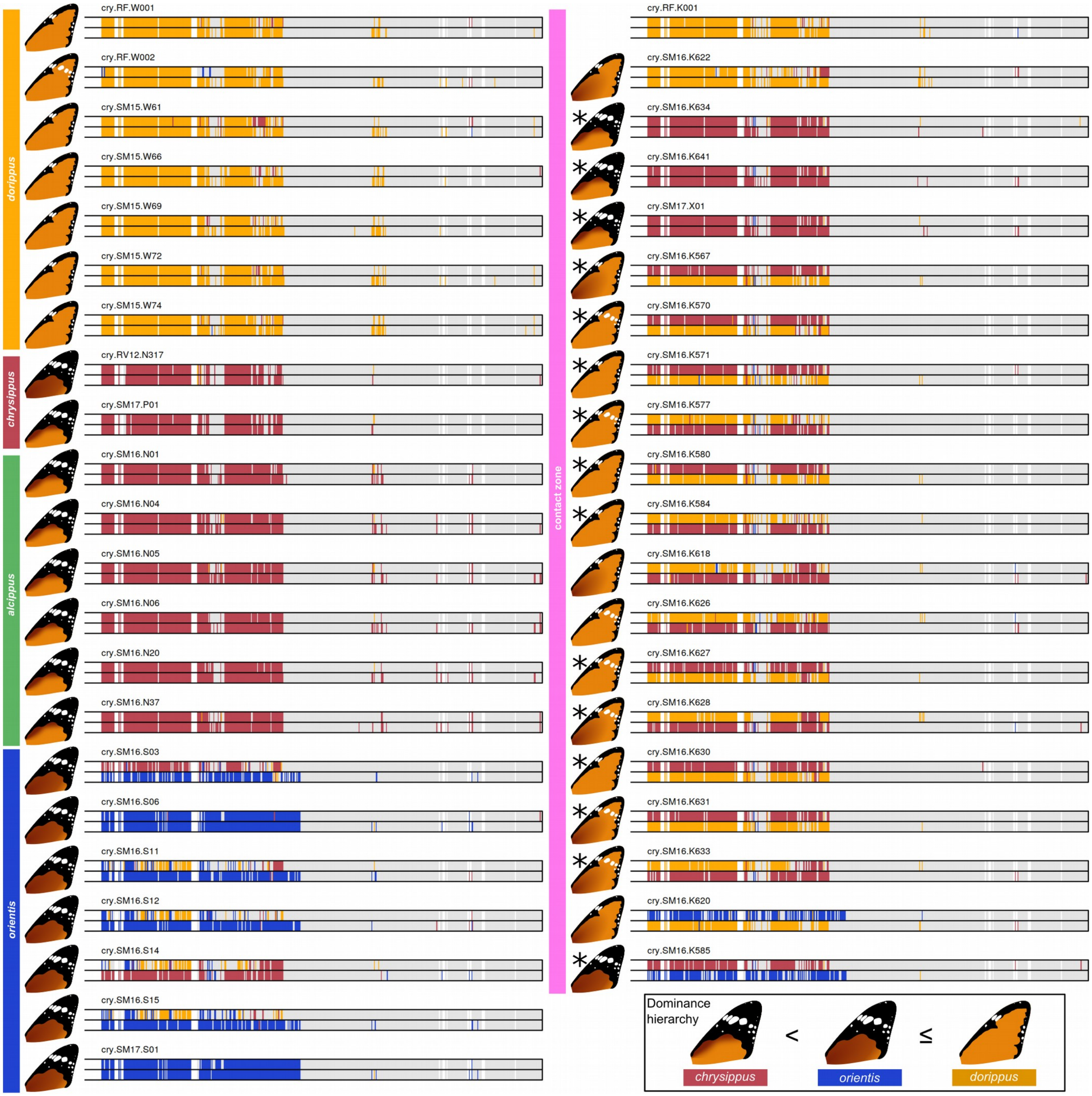


**S4 Fig. Allelic clustering across chromosome 15 for all samples.** Coloured blocks indicate 20 kb windows in which sequence haplotypes could be clustered into one of three genetic clusters (yellow: *dorippus,* red: *chrysippus*, blue: *orientis*) based on pairwise genetic distances (see Methods for details). Windows in grey show insufficient relative divergence to be assigned to a cluster. White gaps indicate missing data. There are three clearly distinct alleles that correspond largely with colour pattern. Heterozygotes and indicate a dominance hierarchy: The $BC_{dorippus}$ allele (yellow) is the most dominant, and produces the *dorippus* phenotype (no black forewing tip). Around half of the heterozygotes with one copy of the *dorippus* allele express the *transiens* phenotype, with white marks on the forewing. The $BC_{orientis}$ allele (blue) corresponds with the *orientis* phenotype (black wing tip and dark background colour). It is dominant over the $BC_{chrysippus}$ allele, which produces the *chysippus* phenotype (black wing tip with light background colour) only when homozygous. There is evidence of recombination in the form of mosaic haplotypes. Finally, samples found to be carrying the neo-W chromosome (see main text) are indicated with an asterisk. All carry the *chrysippus* allele. Note that no phenotype was recorded for the reference genome individual RF.K001.

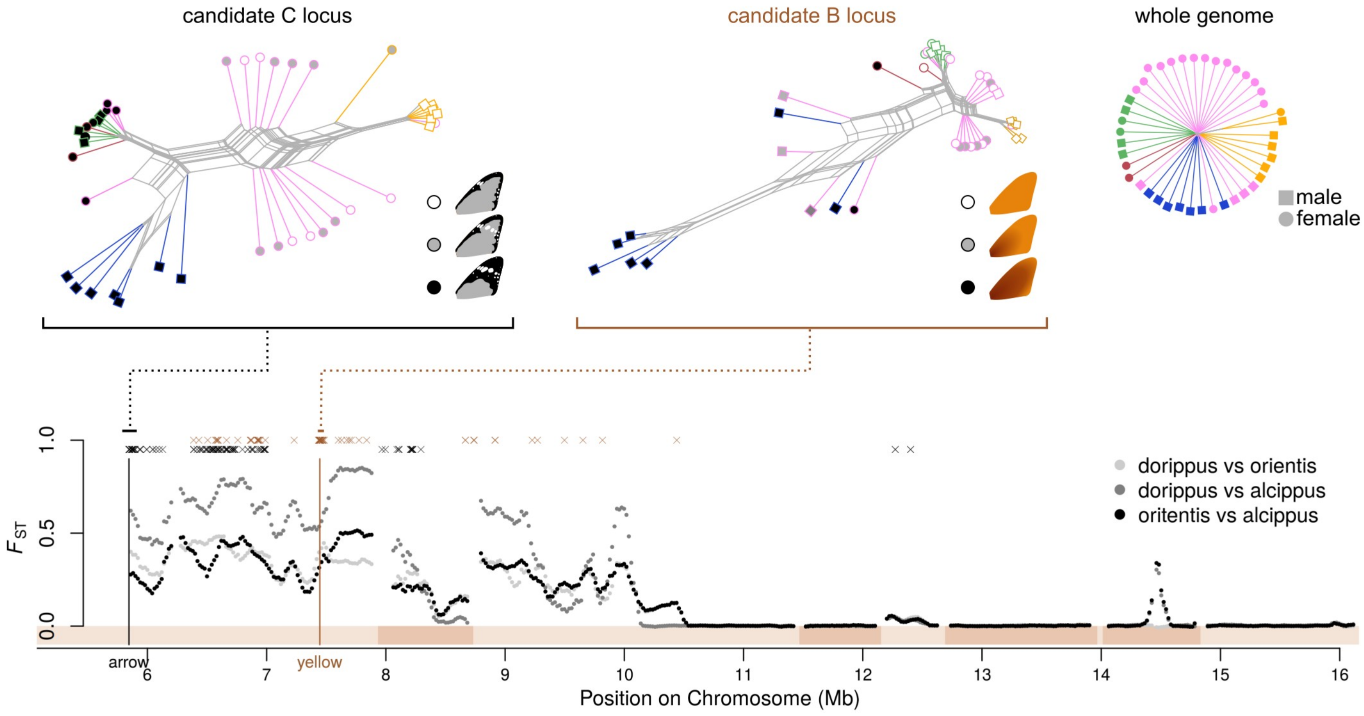


**S5 Fig. Candidate loci for forewing colour pattern on chromosome 15.** Differentiation ($F_{ST}$) is plotted across part of chromosome 15 (bottom). Above the plot, locations of SNPs most strongly associated with the B and C loci (Wald test, 99.99% quantile) are shown: 'B locus' (controlling brown/orange background) in brown and 'C locus' (controlling forwing black tip) in black. The best respective candidate genes *yellow* and *arrow,* are indicated on the plot. At the top, distance-based phylogenetic networks constructed for regions around the candidate genes (30 kb around *yellow* and 100 kb around *arrow*) are shown. Colours indicate subspecies as in Fig. 1A, and shapes indicate sex. Phenotypes for are coded black and white for putative homozygotes and grey for putative heterozygotes. A corresponding network for the whole genome is included for comparison, showing how undifferentiated the subspecies are in general.

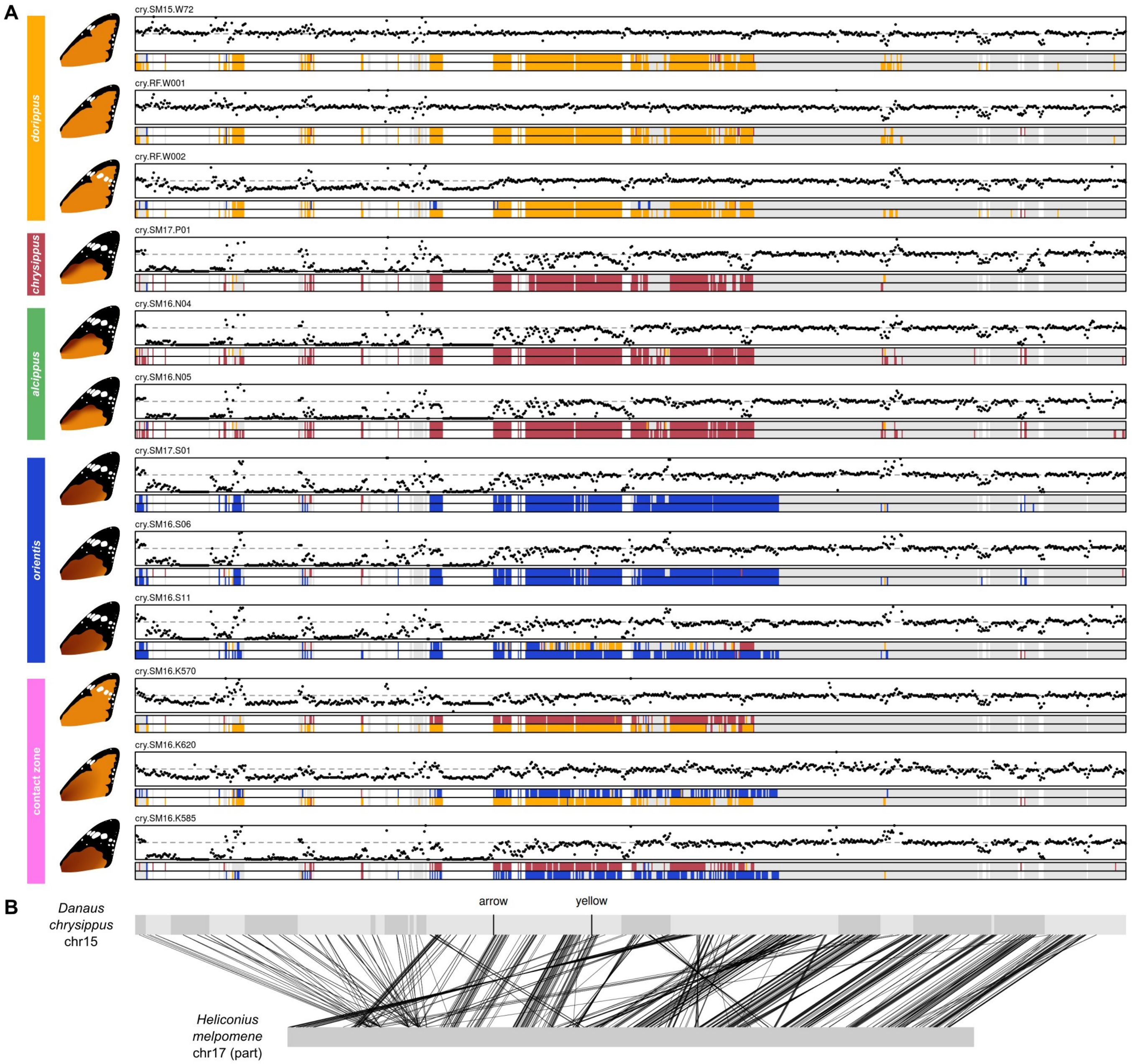


**S6 Fig. Variable coverage reveals a large expansion on chromosome 15.** (**A**) Dots indicate median read coverage in 20 kb windows across chr15, normalised relative to the genome-wide mean (dashed line). Twelve representative individuals are shown. All individuals fall into one of three categories: normal coverage, ~half coverage, or ~zero coverage across the first third (5.84 Mb) of the chromosome, indicating an insertion polymorphism that is either homozygous present/absent or heterozygous. Coloured blocks indicate allelic clustering for each 20 kb window (see S4 Fig), with white indicating gaps in the alignment due to variable sequence coverage. (**B**) Comparison of homologous genes in the *H. melpomene* genome indicates several genes near the proximal end of the chromosome that are duplicated multiple times in our *D. chrysippus* reference genome. Locations of the candidate B and C genes *yellow* and *arrow* (see S5 Fig) are indicated. Scaffolds in the *D. chrysippus* pseudo-chromosomal assembly are alternately shaded light and dark.

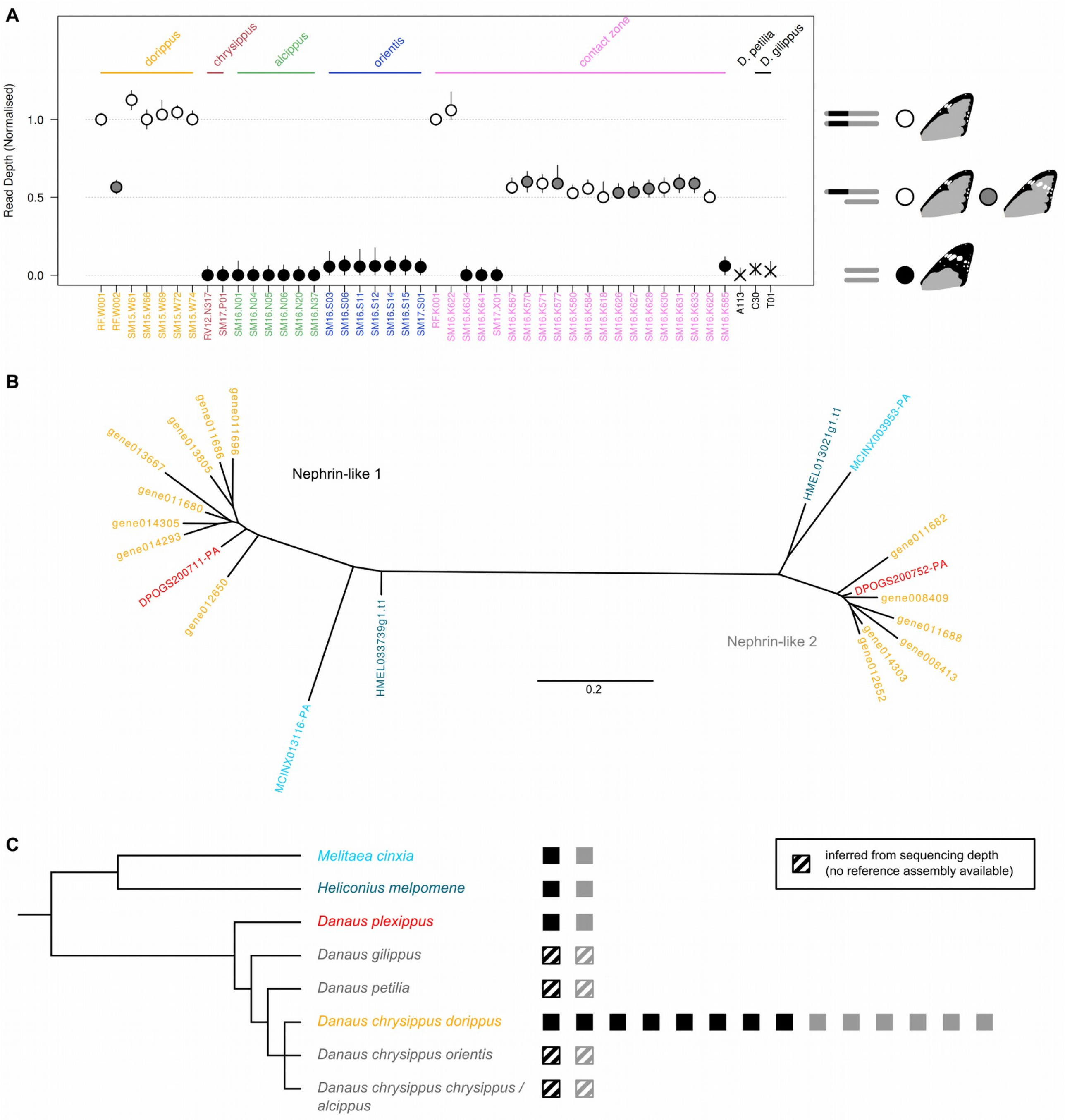


**S7 Fig. An expansion in the $BC^{dorippus}$ allele of chr15 involves multiple gene duplications.** (**A**) Depth of coverage across the expansion region (see S6 Fig), in each individual, normalised by the genome average. Points represent the median coverage over 20 kb windows, and vertical lines indicate the 25% and 75% quantiles. Homozygous individuals with two copies of the expansion have normal depth ~1, heterozygous individuals have depth ~0.5 and those homozygous for a lack of the expansion have depth ~0. There is perfect correspondence between presence of the expansion and the *dorippus* phenotyp (lack of black forewing tip). Heterozygous either the *dorippus* pattern or the "*transiens*" pattern, with white marks on the forewing, consistent with ~50% penetrance described in previous crosses [46]. (**B**) Maximum likelihood phylogeny of Nephrin-like protein sequences encoded by two genes located within the expansion region. Homologous genes from *Danaus plexippus*, *Heliconius melpomene* and *Melitaea cinxia* are included. The tree indicates that the ancestral state in the Nymphalidae is to have two copies of the gene, while the *D. chrysippus* assembly has 14 copies (8 and 6, respectively). (**C**) The number of copies of *nephrin-like 1* and *2* is indicated in black and grey, respectively. Although we have just one assembly from a *D. chrysippus dorippus* sample, the read depth data (see panel A and S6 Fig) suggest that the other *D. chrysippus* subspecies have the ancestral state, lacking the additional copies, as do the two outgroup species: *D. petilia* and *D. gilippus*.

A

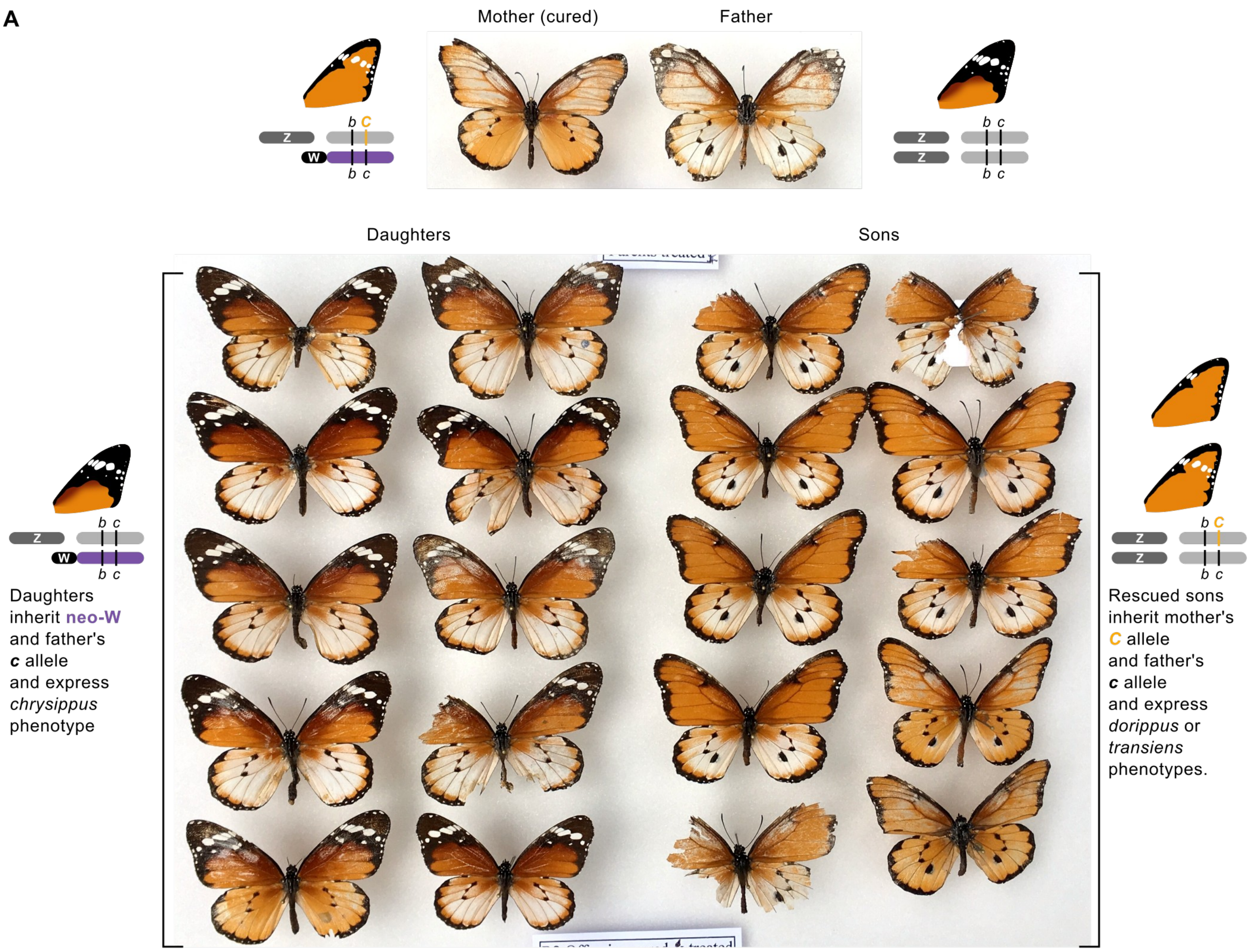


B

| Marker | Mother | Father | Daughters | | | | | | | | | | | | | | | | Sons | | | | | | |
|---|---|---|---|---|---|---|---|---|---|---|---|---|---|---|---|---|---|---|---|---|---|---|---|---|---|
| | | | 1 | 2 | 3 | 4 | 5 | 6 | 7 | 8 | 9 | 10 | 11 | 12 | 13 | 14 | 15 | 16 | 1 | 2 | 3 | 4 | 5 | 6 | 7 |
| sc11_PRC_RFLP_2 ('P') | *Pp* | *pp* | *Pp* | *Pp* | *Pp* | *Pp* | *Pp* | *Pp* | *Pp* | *Pp* | *Pp* | *Pp* | *Pp* | *Pp* | *Pp* | *Pp* | *Pp* | *Pp* | - | *pp* | *pp* | *pp* | *pp* | *pp* | *pp* |
| sc120_PCR_RFLP_2 ('Q') | *qq* | *Qq* | *Qq* | - | *qq* | *qq* | *Qq* | *Qq* | *Qq* | *Qq* | *Qq* | *qq* | *qq* | *Qq* | *Qq* | *qq* | *Qq* | *qq* | *qq* | *qq* | *Qq* | *qq* | *Qq* | *qq* | *qq* |

**S8 Fig. Sex linked inheritance of colour pattern and chr15 in a cured line.** (**A**) Sex linkage of forewing pattern controlled by the BC supergene. A female descending from the contact zone (top left) was cured of *Spiroplasma*. Her *transiens* phenotype indicated that she was heterozygous *Cc* (Fig. 1C). She was crossed with a *cc* male (black forewing tips). Male offsprings (right) who would ordinarily have been killed by *Spiroplasma*, expressed the *dorippus* (or *transiens*) phenotype without black forewing tips, indicating that they had all inherited the *C* allele from their mother. (Note that males can be identified by the additional large black spot on the hindwing). Female offspring (left) all expressed the *chrysippus* phenotype, indicating that they had inherited the recessive *c* allele from both parents. (**B**) Inheritance of two chr15 PCR markers (here designated P and Q) was tracked in the F5 brood of the cured line. One marker (P) was heterozygous in the mother and showed complete sex linkage. The other marker(Q) was heterozygous in the father and segregated independently of sex. These results are consistent with chr15 forming a neo-W in the mother, while both copies of the father's chr15 are autosomal.

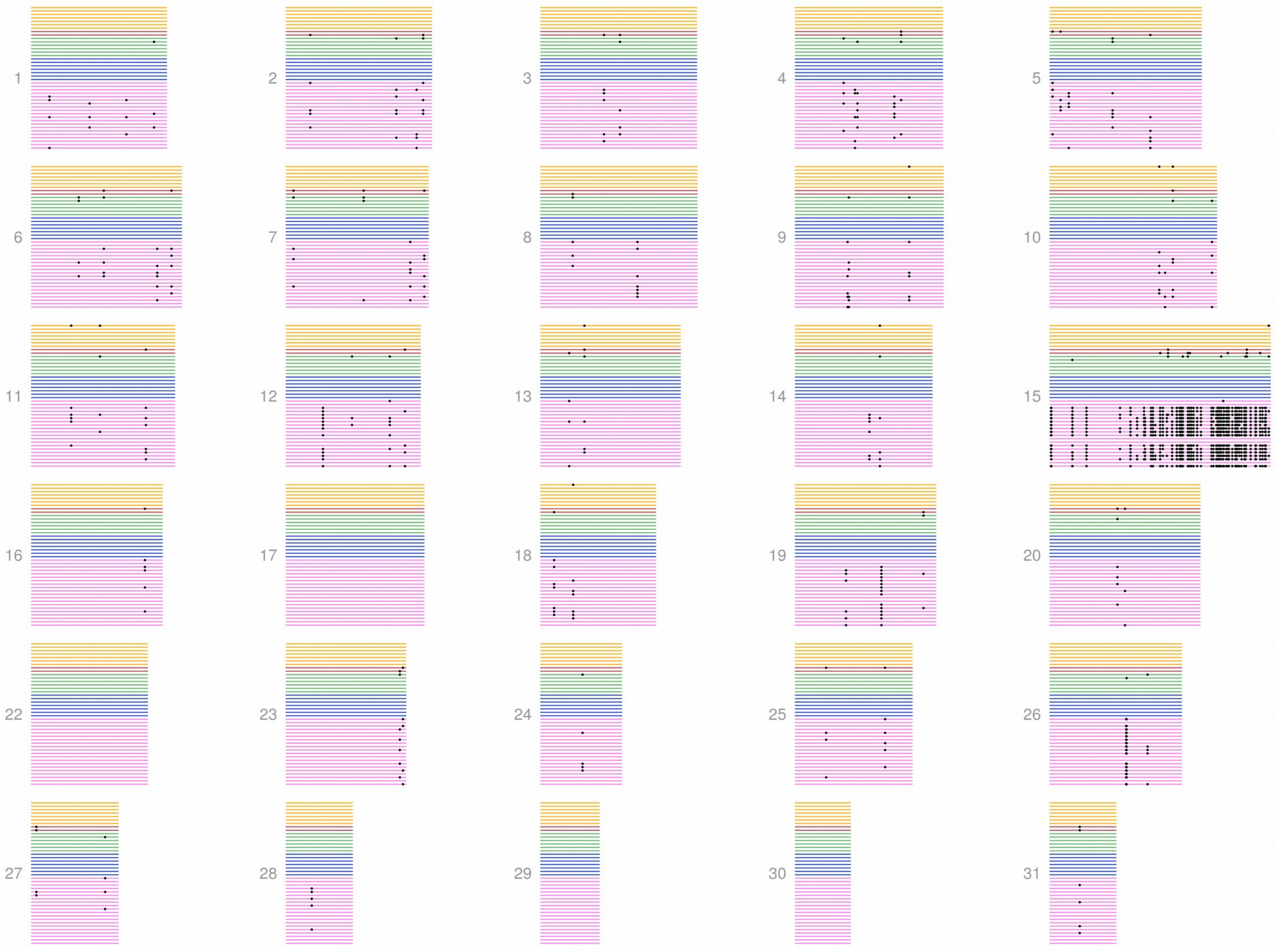


**S9 Fig. Distribution of female-specific mutations identifies the neo-W lineage.** The 30 chromosomes are shown with each line representing an individual, coloured according to population: yellow=*dorippus*, red=*chrysippus*, green=*alcippus*, blue=*orientis*, pink=contact zone. Black points indicate the location of mutations shared by at least four females and absent from males. These are strongly clustered on chromosome 15 (chr15) and shared by a group of contact zone females, indicating that a conserved neo-W haplotype is shared by this female lineage. The noticeable absence of mutations on the proximal (left) region of chr15 reflects the large sequencing gaps corresponding to the expansion cluster in the *dorippus* allele (see S6 Fig).

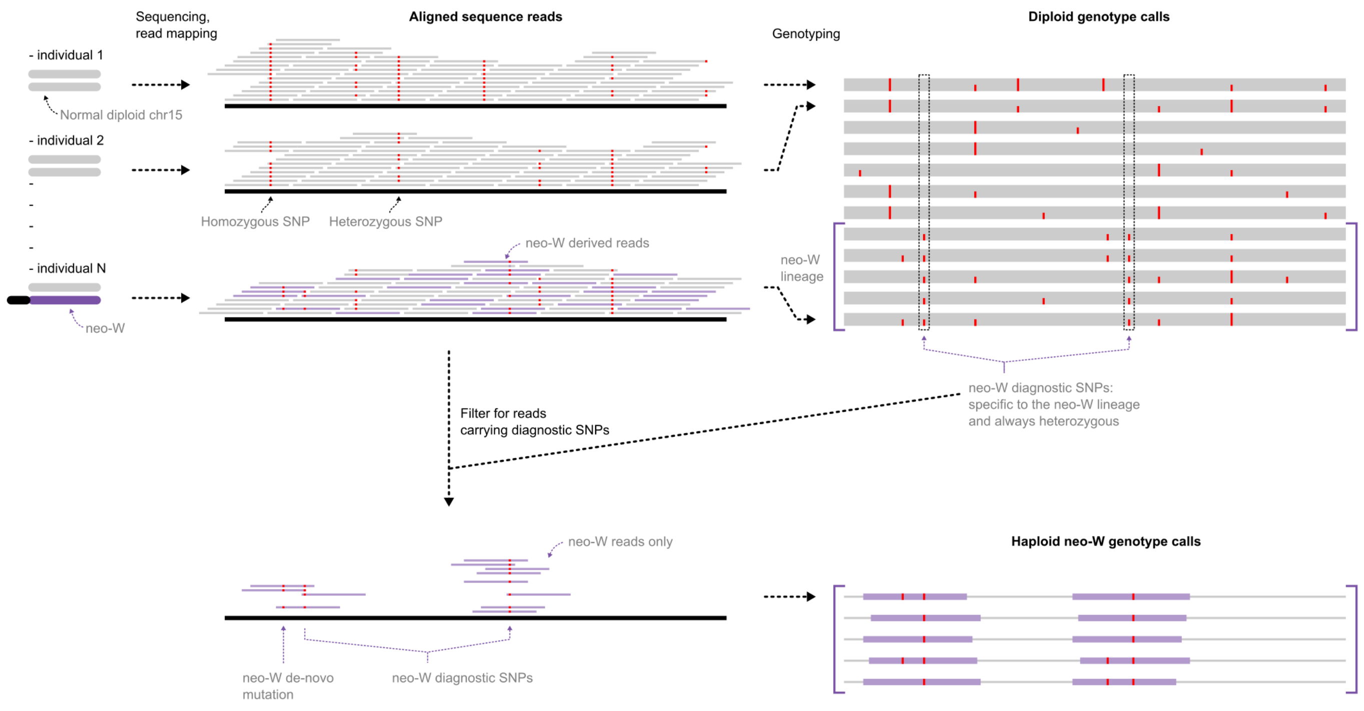


**S10 Fig. Identification of mutations and sequence reads specific to the neo-W.** Schematic representation of the bioinformatic pipeline to isolate the neo-W haplotype from unphased resequencing data. Due to the recency of its formation, sequencing reads from the neo-W are not significantly divergent and will therefore map to the reference genome chr15. The challenge is to separate reads that derive from the neo-W and autosomal haplotypes, despite them all mapping to the same parts of the reference genome. Our solution is to use diagnostic mutations that are unique to the neo-W haplotype and shared by the multiple individuals that carry the neo-W. We identified candidate mutations specific to the neo-W haplotype as those at which all 15 females in the neo-W lineage are heterozygous, while all 27 remaining individuals are homozygous. We then used these candidate neo-W specific mutations to extract sequence reads that are specific to the neo-W. These represent only a fraction of the chromosome, because they represent only the reads carrying diagnostic mutations and their paired-end partners. The identification of these neo-W specific reads allows the identification of additional mutations on the same read that occurred after the formation of the neo-W. These can be used to estimate genetic diversity across the neo-W (accounting for the large amount of missing data), and also to infer a genealogy for the neo-W.

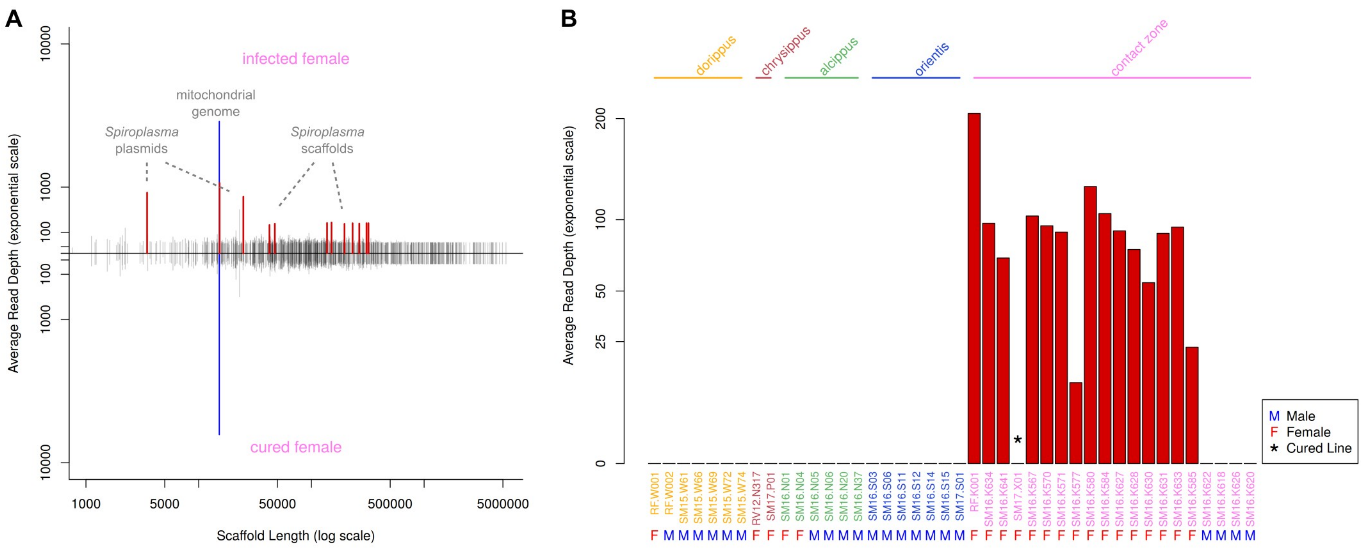


**S11 Fig. Identification of *Spiroplasma* genome and infection status based on read depth.** (**A**) Sequencing read depth of coverage averaged by scaffold (y-axis, exponential scale) and plotted against scaffold length(x-axis, log scale). Depth is shown for a suspected infected female above and a female from the tetracycline-treated 'cured line' below. Scaffolds identified as belonging to the *Spiroplasma* genome are shown in red. The mitochondrial genome is shown in blue. (**B**) Bars show the average depth of reads mapping to the *Spiroplasma* genome for each resequenced *D. chrysippus* individual. Note that all females from the hybrid zone are found to be infected, with the exception of the single individual from the cured line.

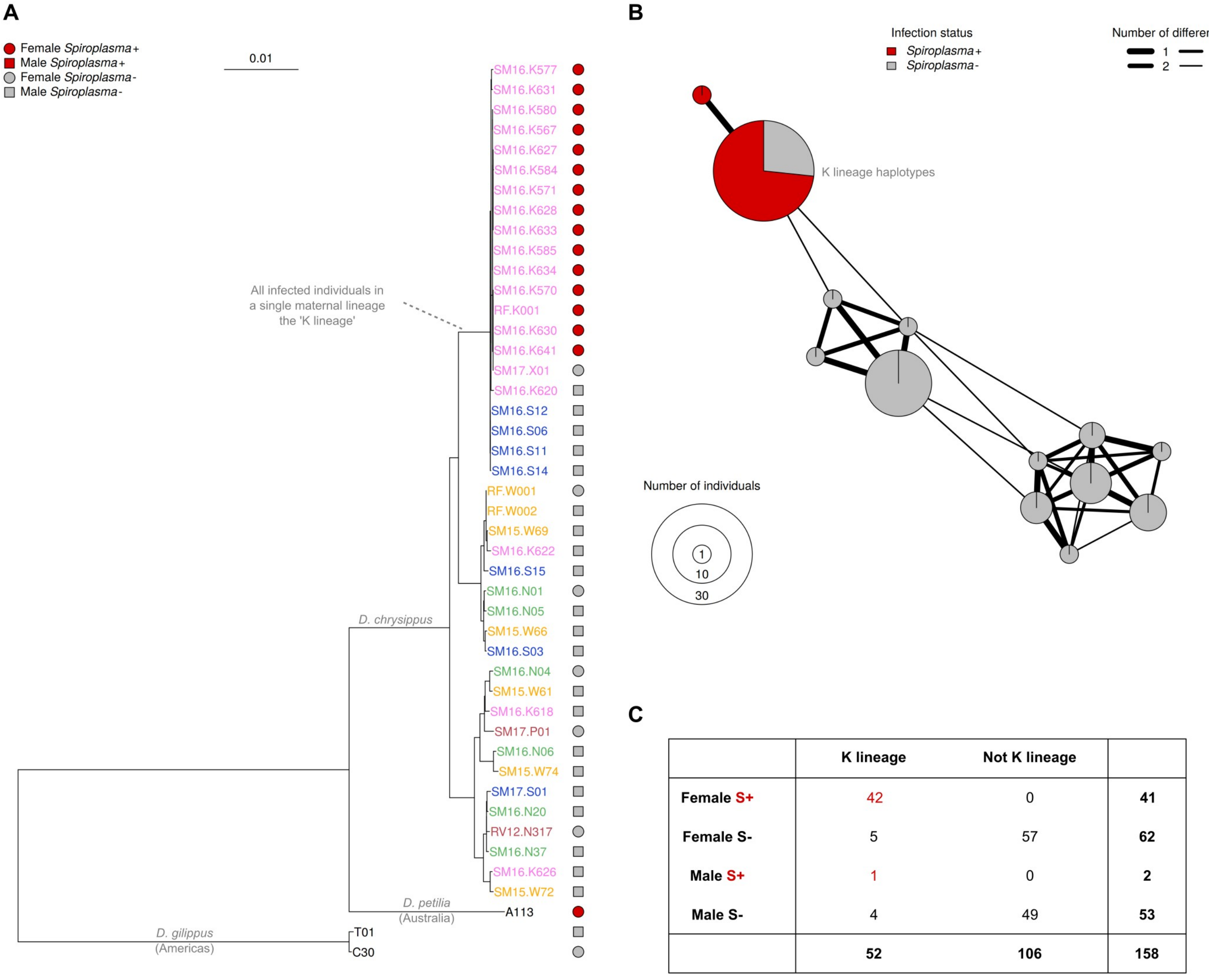


| | K lineage | Not K lineage | |
|---|---|---|---|
| **Female S+** | 42 | 0 | **41** |
| **Female S-** | 5 | 57 | **62** |
| **Male S+** | 1 | 0 | **2** |
| **Male S-** | 4 | 49 | **53** |
| | **52** | **106** | **158** |

**S12 Fig. Association between mitochondrial haplotype and *Spiroplasma* infection.** (**A**) a whole mitochondrial maximum-likelihood phylogeny for the 42 resequenced individuals indicates that all infected *D. chrysippus* females belong to a single mitochondrial clade (here called the K lineage), consistent with strict matrilineal inheritance of *Spiroplasma*. Note that the single *D. petilia* male from Australia was found to be infected by a related *Spiroplasma* strain, but has a different mitochondrial haplotype, indicating an independent infection. (**B**) COI haplotype network for 66 individuals further supports the finding that only K lineage individuals are infected. (**C**) A PCR RFLP for a SNP specific to the K lineage applied to 158 individuals further confirms the finding that only the K lineage carries the infection. Note that one male was found to be infected, probably representing a rare survivor from an infected mother, as has been observed in some experimental crosses [47].

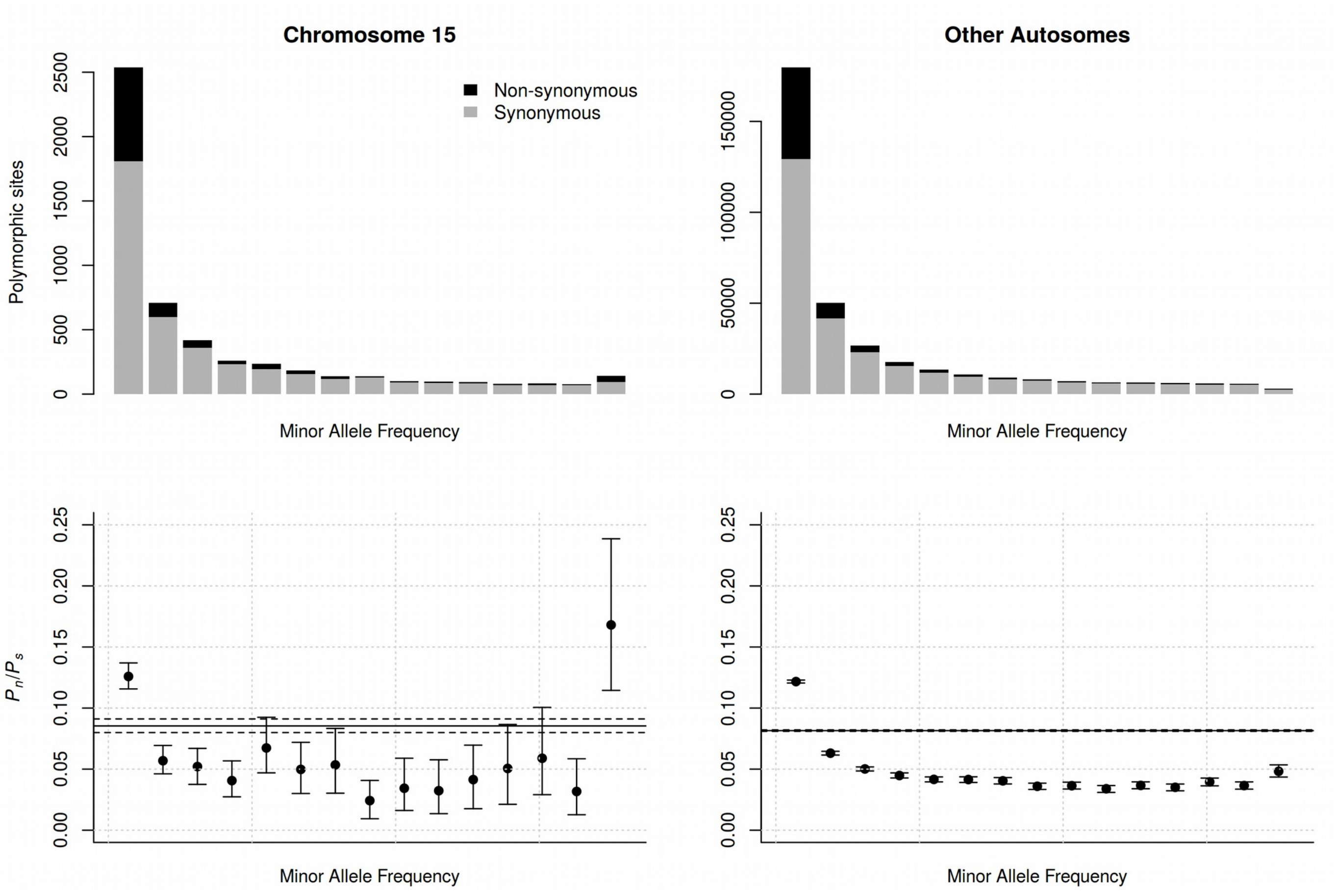


**S13 Fig. Evidence for hitchhiking of non-synonymous mutations on the neo-W.** Barpolots(top) show the frequency distribution of synonymous (grey) and non-synonymous (black) polymorphisms in the neo-W lineage (i.e. contact-zone females carrying the neo-W chromosome). Values for chromosome 15 is shown on the left and combined values across all other autosomes are shown on the right. Below, $P_n/P_s$ (the normalised ratio of non-synonymous to synonymous polymorphisms) is shown for each all frequency class. Error bars show the 95% confidence interval based on 1000 bootstrap replicates. These plots show that non-synonymous polymorphisms are generally skewed toward lower frequency, but that chromosome 15 carries a significant excess of non-synonymous polymorphisms at high frequency in the population. This is consistent with hitchhiking of previously rare mildly-deleterious alleles to high-frequency on the neo-W.

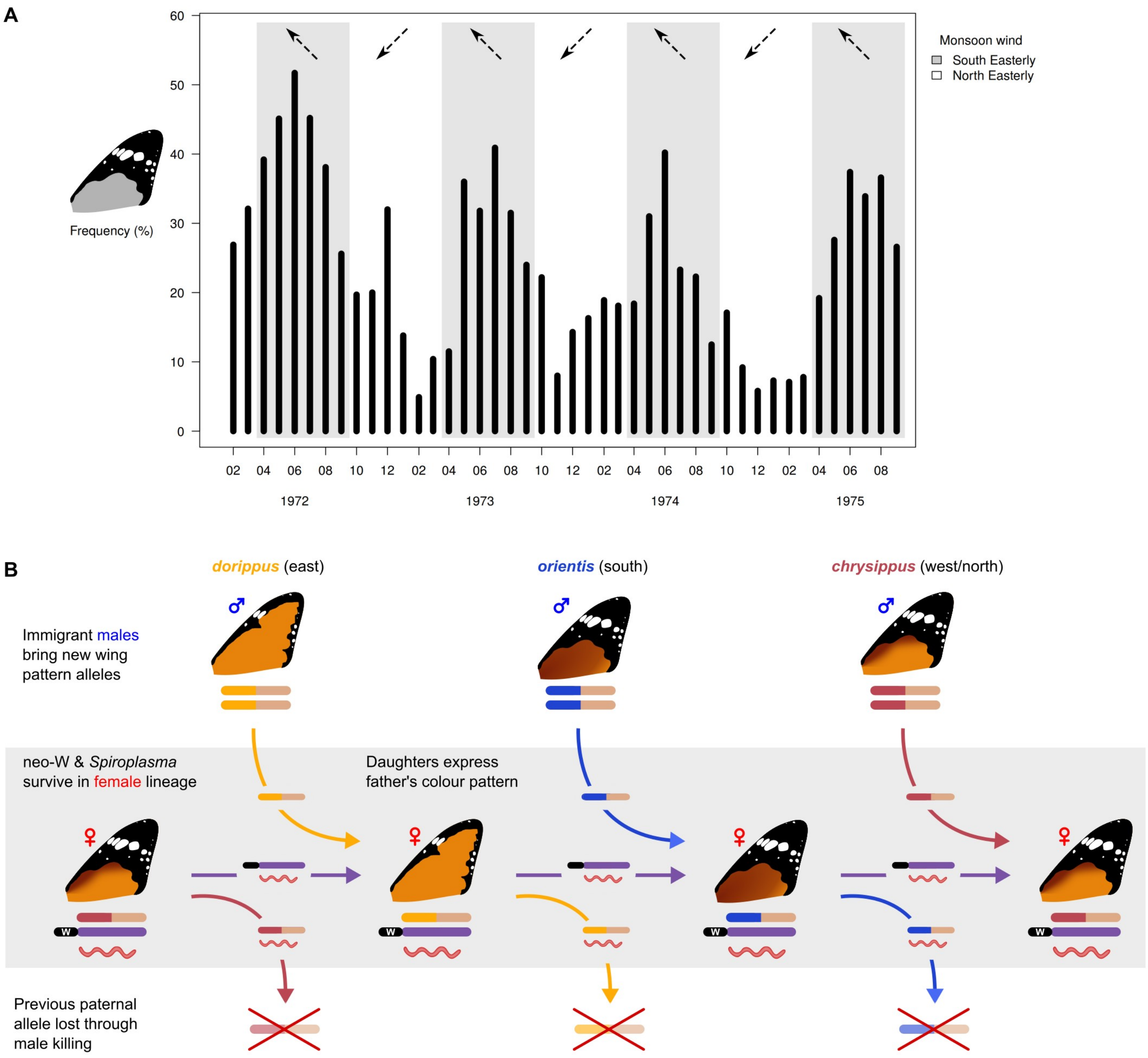


**S14 Fig. Seasonal migration and a genetic sink drive fluctuations in local wing pattern.** (**A**) Average monthly frequencies of the black forewing phenotype (*cc* genotype, *orientis* and *chrysippus* alleles) show how immigration of different subspecies into the contact zone varies seasonally (data from Smith et al. [48], collected at Dar es Salaam between 1972 and 1975). (**B**) Phenotypes of females carrying the neo-W and *Spiroplasma* depend on the source of immigrant males (top row). Each generation, females (middle row) inherit both the neo-W and *Spirplasma* from their mother, and an autosomal chr15 copy from their immigrant father. The neo-W is recessive, causing these females to express their father's phenotype. After persisting in the female for one generation, the autosomal chr15 copy carrying the paternal allele is lost through male killing, i.e. a genetic sink (bottom row). The progression from left to right illustrates how seasonal changes in the predominant source of immigrant males can drive corresponding changes in the phenotypes of the contact zone females.

#### S1 Table. Sequence data used for reference genome assembly

| Library | Type | Read pairs passed (million) | Read pairs trimmed (million) | Mate-pairs | Paired-end | Unknown | Single end | Insert size exp. (Kb) | Insert size obs. (Kb) | Total Gb | Est. depth of cov. |
|---|---|---|---|---|---|---|---|---|---|---|---|
| LIB23244 | PE | 170.64 | 164.84 | | 100% | | | 0.5 | | 82.42 | 329.6 |
| LIB23234 | MP | 18.43 | 18.43 | 70.63% | 17.32% | 6.53% | 5.52% | 11 | 11.5 | 9.21 | 36.85 |
| LIB23235 | MP | 30.88 | 30.87 | 74.71% | 16.35% | 3.88% | 5.07% | 9.5 | 9.5 | 15.44 | 61.74 |
| LIB23236 | MP | 30.36 | 30.35 | 75.50% | 15.91% | 3.48% | 5.11% | 8 | 8.5 | 15.18 | 60.7 |
| LIB23239 | MP | 49.53 | 49.51 | 76.88% | 14.17% | 3.60% | 5.34% | 5 | 4.2 | 24.76 | 99.02 |
| LIB23242 | MP | 42.81 | 42.80 | 76.31% | 15.29% | 4.07% | 4.32% | 2 | 2 | 21.40 | 85.6 |

**S2 Table. Inferred genome properties based on k-mer content**

| | Heterozygosity | Haploid length | Repeat Length | Unique Length | Read Error Rate |
|---|---|---|---|---|---|
| Minimum | 0.0302 | ~249 Mb | 105.6 Mb | 143.2 Mb | 0.00151 |
| Maximum | 0.0306 | ~249.2 Mb | 105.7 Mb | 143.4 Mb | 0.00151 |

**S3 Table. Final *D. chrysippus* assembly statistics**

| | |
|---|---|
| Total number of scaffolds | 726 |
| Total length | 322399471 |
| Largest scaffold | 5321723 |
| GC (%) | 30.52 |
| N50 | 628022 |
| N75 | 628022 |
| L50 | 63 |
| L75 | 141 |
| Total number of scaffolds (>= 0 bp) | 726 |
| Total number of scaffolds (>= 1000 bp) | 726 |
| Total number of scaffolds (>= 5000 bp) | 702 |
| Total number of scaffolds (>= 10000 bp) | 679 |
| Total number of scaffolds (>= 25000 bp) | 605 |
| Total number of scaffolds (>= 50000 bp) | 512 |
| Total length (>= 0 bp) | 322399471 |
| Total length (>= 1000 bp) | 322399471 |
| Total length (>= 5000 bp) | 322335789 |
| Total length (>= 10000 bp) | 322161153 |
| Total length (>= 25000 bp) | 320856651 |
| Total length (>= 50000 bp) | 317312449 |
| Number of N's per 100 kbp | 1113.12 |

**S4 Table. Summarized results of the CEGMA analysis based on 248 CEGs**

| Groups | Number of Proteins | Completeness | Total | Average | Percentage of Orthologs |
|---|---|---|---|---|---|
| Complete | 192 | 77.42 | 224 | 1.17 | 13.02 |
| Group 1 | 52 | 78.89 | 58 | 1.12 | 11.54 |
| Group 2 | 42 | 75.00 | 48 | 1.14 | 9.52 |
| Group 3 | 45 | 73.77 | 52 | 1.16 | 13.33 |
| Group 4 | 53 | 81.54 | 66 | 1.25 | 16.98 |
| Partial | 224 | 90.32 | 304 | 1.36 | 28.57 |
| Group 1 | 60 | 90.91 | 75 | 1.25 | 23.33 |
| Group 2 | 47 | 83.93 | 67 | 1.43 | 31.91 |
| Group 3 | 55 | 90.16 | 75 | 1.36 | 29.09 |
| Group 4 | 62 | 95.38 | 87 | 1.40 | 30.65 |

**S5 Table. BUSCO statistics for 3 clades**

| Clade | Total BUSCO | Complete BUSCO | Single copy | Duplicated | Fragmented | Missing |
|---|---|---|---|---|---|---|
| Eukaryota | 303 | 279 | 273 | 6 | 4 | 20 |
| Arthropoda | 1066 | 1002 | 994 | 8 | 14 | 50 |
| Insecta | 1658 | 1563 | 1547 | 16 | 17 | 78 |

**S6 Table. Summary of gene features in *D. chrysippus* genome**

| | |
|---|---|
| Total sequence length | 323361855 |
| Number of genes | 16654 |
| Number of mRNAs | 18300 |
| Number of exons | 98865 |
| Number of introns | 80565 |
| Number of CDS | 15682 |
| Total gene length | 63146110 |
| Total mRNA length | 66260632 |
| Total exon length | 20536415 |
| Total intron length | 45885347 |
| Total CDS length | 20328882 |
| mean gene length | 3792 |
| mean mRNA length | 3621 |
| mean exon length | 208 |
| mean intron length | 570 |
| mean CDS length | 1296 |
| % of genome covered by genes | 19.5% |
| % of genome covered by CDS | 6.3% |

**S7 Table. Orthogroups summary statistics**

| | |
|---|---|
| Number of genes | 74925 |
| Number of genes in orthogroups | 66709 |
| Number of unassigned genes | 8216 |
| Percentage of genes in orthogroups | 89.0 |
| Percentage of unassigned genes | 11.0 |
| Number of orthogroups | 12738 |
| Number of species-specific orthogroups | 39 |
| Number of genes in species-specific orthogroups | 173 |
| Percentage of genes in species-specific orthogroups | 0.2 |
| Mean orthogroup size | 5.2 |
| Median orthogroup size | 5.0 |
| G50 (assigned genes) | 5 |
| G50 (all genes) | 5 |
| O50 (assigned genes) | 4393 |
| O50 (all genes) | 5214 |
| Number of orthogroups with all species present | 7355 |
| Number of single-copy orthogroups | 4858 |

**S8 Table. Distribution of orthogroups in different species**

| Species | #genes | #genes in OG | #OG containing species |
|---|---|---|---|
| *Bombyx mori* | 14623 | 12653 (86.5%) | 10211 (80.2%) |
| *Danaus chrysippus* | 15675 | 14895 (95%) | 11343 (89%) |
| *Danaus plexippus* | 15130 | 14241 (94.1%) | 12015 (94.3%) |
| *Heliconius melpomene* | 12829 | 11954 (93.2%) | 10034 (78.8%) |
| *Melitaea cincia* | 16668 | 12966 (77.8%) | 10379 (81.5%) |

**S9 Table. Sample information for population genomic analyses**

| ID | Taxon | Sex | Location | Lat. | Long. | Total Gb | Mean Depth | Accession |
|---|---|---|---|---|---|---|---|---|
| RF.W001 | *Danaus chrysippus dorippus* | F | Watamu, Kenya | -3.33 | 40.02 | 11.49* | 35.94* | |
| RF.W002 | *Danaus chrysippus dorippus* | M | Watamu, Kenya | -3.33 | 40.02 | 10.38* | 32.45* | |
| SM15.W61 | *Danaus chrysippus dorippus* | M | Watamu, Kenya | -3.33 | 40.02 | 7.47 | 23.37 | |
| SM15.W66 | *Danaus chrysippus dorippus* | M | Watamu, Kenya | -3.33 | 40.02 | 7.15 | 22.35 | |
| SM15.W69 | *Danaus chrysippus dorippus* | M | Watamu, Kenya | -3.33 | 40.02 | 6.95 | 21.73 | |
| SM15.W72 | *Danaus chrysippus dorippus* | M | Watamu, Kenya | -3.33 | 40.02 | 9.82 | 30.72 | |
| SM15.W74 | *Danaus chrysippus dorippus* | M | Watamu, Kenya | -3.33 | 40.02 | 7.86 | 24.57 | |
| RV12.N317 | *Danaus chrysippus chrysippus* | F | El Haouareb, Tunisia | 35.549 | 9.754 | 7.54 | 23.59 | |
| SM17.P01 | *Danaus chrysippus chrysippus* | F | Philippines (via Stratford Butterfly Farm) | ? | ? | 8.15 | 25.48 | |
| SM16.N01 | *Danaus chrysippus alcippus* | F | Kanyang, Nigeria | 6.24 | 8.97 | 8.56 | 26.76 | |
| SM16.N04 | *Danaus chrysippus alcippus* | F | Kanyang, Nigeria | 6.24 | 8.97 | 7.73 | 24.17 | |
| SM16.N05 | *Danaus chrysippus alcippus* | M | Kanyang, Nigeria | 6.24 | 8.97 | 7.18 | 22.45 | |
| SM16.N06 | *Danaus chrysippus alcippus* | M | Kanyang, Nigeria | 6.24 | 8.97 | 6.84 | 21.4 | |
| SM16.N20 | *Danaus chrysippus alcippus* | M | Kanyang, Nigeria | 6.24 | 8.97 | 8.11 | 25.36 | |
| SM16.N37 | *Danaus chrysippus alcippus* | M | Kanyang, Nigeria | 6.24 | 8.97 | 8.11 | 25.37 | |
| SM16.S03 | *Danaus chrysippus orientis* | M | Magaliesberg, South Africa | -26.02 | 27.51 | 8.39 | 26.23 | |
| SM16.S06 | *Danaus chrysippus orientis* | M | Magaliesberg, South Africa | -26.02 | 27.51 | 7.12 | 22.28 | |
| SM16.S11 | *Danaus chrysippus orientis* | M | Magaliesberg, South Africa | -26.02 | 27.51 | 7.91 | 24.72 | |
| SM16.S12 | *Danaus chrysippus orientis* | M | Magaliesberg, South Africa | -26.02 | 27.51 | 7.32 | 22.9 | |
| SM16.S14 | *Danaus chrysippus orientis* | M | Magaliesberg, South Africa | -26.02 | 27.51 | 7.58 | 23.7 | |
| SM16.S15 | *Danaus chrysippus orientis* | M | Magaliesberg, South Africa | -26.02 | 27.51 | 7.69 | 24.04 | |
| SM17.S01 | *Danaus chrysippus orientis* | M | Magaliesberg, South Africa | -26.02 | 27.51 | 8.30 | 25.97 | |
| RF.K001 | contact zone | F | Nairobi, Kenya | -1.39 | 36.82 | 11.70* | 36.59* | |

| | | | | | | | |
|---|---|---|---|---|---|---|---|
| SM16.K622 | contact zone | M | Nairobi, Kenya | -1.39 | 36.82 | 7.72 | 24.15 |
| SM16.K634 | contact zone | F | Nairobi, Kenya | -1.39 | 36.82 | 7.72 | 24.13 |
| SM16.K641 | contact zone | F | Nairobi, Kenya | -1.39 | 36.82 | 9.12 | 28.53 |
| SM16.K567 | contact zone | F | Nairobi, Kenya | -1.39 | 36.82 | 7.30 | 22.83 |
| SM16.K570 | contact zone | F | Nairobi, Kenya | -1.39 | 36.82 | 6.74 | 21.07 |
| SM16.K571 | contact zone | F | Nairobi, Kenya | -1.39 | 36.82 | 7.64 | 23.89 |
| SM16.K577 | contact zone | F | Nairobi, Kenya | -1.39 | 36.82 | 8.20 | 25.63 |
| SM16.K580 | contact zone | F | Nairobi, Kenya | -1.39 | 36.82 | 8.28 | 25.9 |
| SM16.K584 | contact zone | F | Nairobi, Kenya | -1.39 | 36.82 | 8.15 | 25.5 |
| SM16.K618 | contact zone | M | Nairobi, Kenya | -1.39 | 36.82 | 8.61 | 26.94 |
| SM16.K626 | contact zone | M | Nairobi, Kenya | -1.39 | 36.82 | 7.63 | 23.87 |
| SM16.K627 | contact zone | F | Nairobi, Kenya | -1.39 | 36.82 | 6.67 | 20.87 |
| SM16.K628 | contact zone | F | Nairobi, Kenya | -1.39 | 36.82 | 7.85 | 24.55 |
| SM16.K630 | contact zone | F | Nairobi, Kenya | -1.39 | 36.82 | 7.20 | 22.53 |
| SM16.K631 | contact zone | F | Nairobi, Kenya | -1.39 | 36.82 | 7.61 | 23.8 |
| SM16.K633 | contact zone | F | Nairobi, Kenya | -1.39 | 36.82 | 7.59 | 23.75 |
| SM16.K620 | contact zone | M | Nairobi, Kenya | -1.39 | 36.82 | 8.79 | 27.49 |
| SM16.K585 | contact zone | F | Nairobi, Kenya | -1.39 | 36.82 | 7.38 | 23.07 |
| SM17.X01 | cured line | F | Stock | NA | NA | 8.32 | 26.02 |
| pet.A113 | *Danaus petilia* | M | Australia | ? | ? | 7.37 | 23.05 |
| gil.C30 | *Danaus gilippus* | F | Costa Rica | ? | ? | 8.00 | 25.02 |
| gil.T01 | *Danaus gilippus* | M | Texas, USA | ? | ? | 15.41 | 48.19 |

* indicates sequence content after down-sampling to reduce file size and processing time.

**S10 Table. Closest genes to SNPs most strongly associated with colour pattern traits.** Shading indicates the best candidate gene(s) with the most nearby associated SNPs.

| Trait Association | Gene | Nearby SNPs | Description | e-Value |
|---|---|---|---|---|
| A | gene015410 | 3 | transmembrane and TPR repeat-containing protein CG4341-like | 9.58843E-151 |
| A | gene015308 | 2 | pheromone biosynthesis-activating neuropeptide receptor isoform B | 1.0891E-33 |
| A | gene003441 | 1 | putative Calnexin | 0 |
| A | gene003471 | 1 | transcriptional enhancer factor TEF-1 like protein | 0 |
| A | gene005973 | 1 | serine protease 5 | 1.9592E-154 |
| A | gene006029 | 1 | hypothetical protein KGM_208582 | 0 |
| A | gene006044 | 1 | REPAT32 protein | 1.8188E-28 |
| A | gene006082 | 1 | uncharacterized protein LOC113497409 | 1.74006E-133 |
| A | gene013049 | 1 | hypothetical protein KGM_211895 | 2.11066E-34 |
| A | gene013615 | 1 | splicing factor proline- and glutamine-rich | 0 |
| A | gene015600 | 1 | cuticle protein | 4.02653E-118 |
| B | gene001666 | 4 | yellow | 0 |
| B | gene001484 | 1 | hypothetical protein KGM_206584 | 1.96903E-104 |
| B | gene003564 | 1 | amino acid transporter | 0 |
| B | gene005522 | 1 | serine protease inhibitor 33 precursor | 0 |
| B | gene009383 | 1 | UDP-glycosyltransferase UGT33J1 | 0 |
| B | gene009488 | 1 | Ecdysteroid UDP-glucosyltransferase | 4.26885E-143 |
| B | gene009489 | 1 | UDP-glycosyltransferase UGT33F1 | 0 |
| B | gene009492 | 1 | UDP-glycosyltransferase UGT33F1 | 0 |
| B | gene009493 | 1 | UDP-glycosyltransferase UGT33F1 | 0 |
| B | gene009511 | 1 | uncharacterized protein LOC112045604 | 0 |
| B | gene013199 | 1 | hypothetical protein KGM_206574 | 5.47109E-67 |
| B | gene013913 | 1 | putative structural maintenance of chromosomes 5 smc5 | 0 |
| B | gene013915 | 1 | hemolymph proteinase 24 | 1.29126E-164 |
| C | gene000403 | 2 | low-density lipoprotein receptor-related protein 6 (arrow) | 0 |
| C | gene001633 | 2 | hemolin-interacting protein | 1.23689E-84 |
| C | gene001631 | 1 | hypothetical protein KGM_211925 | 1.56351E-117 |
| C | gene003274 | 1 | hypothetical protein KGM_200756 | 1.70394E-09 |
| C | gene003458 | 1 | ribosomal protein L27A | 4.35329E-76 |
| C | gene003477 | 1 | hypothetical protein KGM_205676 | 0 |
| C | gene003558 | 1 | tetraspanin-9 like protein | 5.36757E-139 |
| C | gene011502 | 1 | trypsin protein precursor | 1.43537E-173 |
| C | gene011692 | 1 | uncharacterized protein LOC113397363 | 2.12478E-66 |
| C | gene011696 | 1 | hypothetical protein KGM_200711 | 0 |
| C | gene013034 | 1 | orexin receptor type 1-like | 6.6138E-138 |

**S11 Table. Details of genotyping assays**

| Assay Details | Target | Forward primer | Reverse primer |
|---|---|---|---|
| "sc11_PCR_RFLP_2"<br>(Fragment length polymorphisms without cleavage) | sc0000011<br>571266-571486<br>(chr15) | AAGAGTTTTAGCGCCGTAAG | CAACTCCTTGTCGTAATGATGA |
| "sc120_PCR_RFLP_2"<br>RFLP with enzyme NdeI | sc0000120<br>387153-387517<br>(chr15) | CTTTCGCAAAGCCAAGGGAC | TAATCGAGGCGACCACAGTG |
| "COI2"<br>Sequencing and RFLP with enzyme BsmI | Cytochrome Oxidase Subunit I (COI), Mitochondirial | GCTTAAACTCAGCCATTTTATTAGCG | TGGATCTCCTCCTCCAGCAG |
| "GDP1"<br>PCR assay for *Spiroplasma* infection | *Spiroplasma* glycero-phosphoryl diester phospho-diesterase (GDP) gene | GAAAATTTGCCAAGCAGTAGAG | AACTACGGAAATTGAAGGATGC |

**S12 Table. Mitochondrial haplotype and infection status of 158 samples screened**

Screening for mitochondrial type was either through direct sequencing or PCR-RFLP for a diagnostic SNP in the COI amplicon. Screening for infection status was either based on resequecning data (see S11 Fig) or by PCR amplification of the *Spiroplasma* GDP1 gene.

| ID | Sex | wild/reared | latitude | longitude | K lineage SNP | Infection Status (GDP PCR) | PCR or Whole Genome Sequencing | COI Seqenced |
|---|---|---|---|---|---|---|---|---|
| SM16.S03 | M | wild | -26.02 | 27.51 | 0 | 0 | WGS | yes |
| SM16.S06 | M | wild | -26.02 | 27.51 | 0 | 0 | WGS | yes |
| SM16.S11 | M | wild | -26.02 | 27.51 | 0 | 0 | WGS | yes |
| SM16.S12 | M | wild | -26.02 | 27.51 | 0 | 0 | WGS | yes |
| SM16.S14 | M | wild | -26.02 | 27.51 | 0 | 0 | WGS | yes |
| SM16.S15 | M | wild | -26.02 | 27.51 | 0 | 0 | WGS | yes |
| SM17.S01 | M | wild | -26.02 | 27.51 | 0 | 0 | WGS | yes |
| SM17.H01 | F | wild | -5.7175 | -15.9249 | 0 | 0 | PCR | no |
| SM17.H02 | F | wild | -5.7175 | -15.9249 | 0 | 0 | PCR | no |
| SM17.H03 | F | wild | -5.7175 | -15.9249 | 0 | 0 | PCR | no |
| SM17.H04 | M | wild | -5.7175 | -15.9249 | 0 | 0 | PCR | no |
| DS06.A07 | F | wild | -5.1 | 38.63 | 0 | 0 | PCR | no |
| DS06.A12 | F | wild | -5.1 | 38.63 | 0 | 0 | PCR | no |
| DS06.A13 | F | wild | -5.1 | 38.63 | 0 | 0 | PCR | no |
| DS06.A14 | F | wild | -5.1 | 38.63 | 0 | 0 | PCR | no |
| DS06.A16 | F | wild | -5.1 | 38.63 | 0 | 0 | PCR | no |
| DS06.A17 | F | wild | -5.1 | 38.63 | 0 | 0 | PCR | no |
| DS06.A18 | F | wild | -5.1 | 38.63 | 0 | 0 | PCR | no |
| DS06.A19 | F | wild | -5.1 | 38.63 | 0 | 0 | PCR | no |
| RF.W001 | F | wild | -3.33 | 40.02 | 0 | 0 | WGS | yes |
| RF.W002 | M | wild | -3.33 | 40.02 | 0 | 0 | WGS | yes |
| SM15.W61 | M | wild | -3.33 | 40.02 | 0 | 0 | WGS | yes |
| SM15.W66 | M | wild | -3.33 | 40.02 | 0 | 0 | WGS | yes |
| SM15.W69 | M | wild | -3.33 | 40.02 | 0 | 0 | WGS | yes |
| SM15.W72 | M | wild | -3.33 | 40.02 | 0 | 0 | WGS | yes |
| SM15.W74 | M | wild | -3.33 | 40.02 | 0 | 0 | WGS | yes |
| SM16.W73 | M | wild | -3.33 | 40.02 | 0 | 0 | PCR | no |
| SM17.W01 | F | wild | -3.33 | 40.02 | 0 | 0 | PCR | no |
| SM17.R20 | F | wild | -2.22 | 30.12 | 0 | 0 | PCR | no |

| | | | | | | | | |
|---|---|---|---|---|---|---|---|---|
| SM17.R21 | M | wild | -2.22 | 30.12 | 0 | 0 | PCR | no |
| SM17.R22 | M | wild | -2.22 | 30.12 | 0 | 0 | PCR | no |
| SM17.R23 | F | wild | -2.22 | 30.12 | 1 | 1 | PCR | yes |
| SM18.R009 | M | wild | -2.22 | 30.12 | 1 | 0 | PCR | no |
| SM18.R010 | F | wild | -2.22 | 30.12 | 0 | 0 | PCR | no |
| SM18.R011 | F | wild | -2.22 | 30.12 | 0 | 0 | PCR | no |
| SM18.R012 | M | wild | -2.22 | 30.12 | 1 | 0 | PCR | yes |
| SM18.M002 | M | wild | -1.9 | 36.048 | 0 | 0 | PCR | no |
| SM18.M003 | F | wild | -1.9 | 36.048 | 0 | 0 | PCR | no |
| RF.K001 | F | wild | -1.39 | 36.82 | 1 | 1 | WGS | yes |
| SM16.K567 | F | wild | -1.39 | 36.82 | 1 | 1 | WGS & PCR | yes |
| SM16.K570 | F | wild | -1.39 | 36.82 | 1 | 1 | WGS | yes |
| SM16.K571 | F | wild | -1.39 | 36.82 | 1 | 1 | WGS & PCR | yes |
| SM16.K577 | F | wild | -1.39 | 36.82 | 1 | 1 | WGS & PCR | yes |
| SM16.K580 | F | wild | -1.39 | 36.82 | 1 | 1 | WGS & PCR | yes |
| SM16.K583 | F | wild | -1.39 | 36.82 | 1 | 1 | PCR | no |
| SM16.K584 | F | wild | -1.39 | 36.82 | 1 | 1 | WGS & PCR | yes |
| SM16.K585 | F | wild | -1.39 | 36.82 | 1 | 1 | WGS & PCR | yes |
| SM16.K618 | M | wild | -1.39 | 36.82 | 0 | 0 | WGS & PCR | yes |
| SM16.K619 | M | wild | -1.39 | 36.82 | 0 | 0 | PCR | no |
| SM16.K620 | M | wild | -1.39 | 36.82 | 1 | 0 | WGS & PCR | yes |
| SM16.K622 | M | wild | -1.39 | 36.82 | 0 | 0 | WGS & PCR | yes |
| SM16.K624 | M | wild | -1.39 | 36.82 | 1 | 1 | PCR | no |
| SM16.K625 | M | wild | -1.39 | 36.82 | 1 | 0 | PCR | no |
| SM16.K626 | M | wild | -1.39 | 36.82 | 0 | 0 | WGS & PCR | yes |
| SM16.K627 | F | wild | -1.39 | 36.82 | 1 | 1 | WGS & PCR | yes |
| SM16.K628 | F | wild | -1.39 | 36.82 | 1 | 1 | WGS & PCR | yes |
| SM16.K630 | F | wild | -1.39 | 36.82 | 1 | 1 | WGS & PCR | yes |
| SM16.K631 | F | wild | -1.39 | 36.82 | 1 | 1 | WGS | yes |
| SM16.K633 | F | wild | -1.39 | 36.82 | 1 | 1 | WGS & PCR | yes |
| SM16.K634 | F | wild | -1.39 | 36.82 | 1 | 1 | WGS & PCR | yes |
| SM16.K641 | F | wild | -1.39 | 36.82 | 1 | 1 | WGS & PCR | yes |
| SM16.K700 | F | wild | -1.39 | 36.82 | 1 | 1 | PCR | no |
| ABRI.2018.3087 | F | wild | -1.304 | 36.693 | 1 | 0 | PCR | no |
| ABRI.2018.3088 | M | wild | -1.304 | 36.693 | 1 | 1 | PCR | no |
| ABRI.2018.3089 | F | wild | -1.304 | 36.693 | 1 | 1 | PCR | no |
| ABRI.2018.3090 | F | wild | -1.304 | 36.693 | 0 | 0 | PCR | yes |
| ABRI.2018.3091 | F | wild | -1.304 | 36.693 | 1 | 1 | PCR | no |
| ABRI.2018.3092 | F | wild | -1.304 | 36.693 | 1 | 1 | PCR | no |

| | | | | | | | | |
|---|---|---|---|---|---|---|---|---|
| ABRI.2018.3093 | F | wild | -1.304 | 36.693 | 1 | 1 | PCR | yes |
| ABRI.2018.3094 | F | wild | -1.304 | 36.693 | 1 | 0 | PCR | yes |
| ABRI.2018.3095 | F | wild | -1.304 | 36.693 | 1 | 0 | PCR | no |
| ABRI.2018.3096 | F | wild | -1.304 | 36.693 | 0 | 0 | PCR | no |
| ABRI.2018.3097 | F | wild | -1.304 | 36.693 | 1 | 1 | PCR | no |
| ABRI.2018.3098 | F | wild | -1.304 | 36.693 | 1 | 0 | PCR | no |
| ABRI.2018.3099 | F | wild | -1.304 | 36.693 | 1 | 1 | PCR | no |
| ABRI.2018.3100 | F | wild | -1.304 | 36.693 | 1 | 1 | PCR | no |
| SM15.K01 | F | wild | -1.304 | 36.693 | 0 | 0 | PCR | yes |
| SM15.K02 | M | wild | -1.304 | 36.693 | 0 | 0 | PCR | yes |
| SM15.K03 | F | wild | -1.304 | 36.693 | 0 | 0 | PCR | yes |
| SM15.K04 | M | wild | -1.304 | 36.693 | 0 | 0 | PCR | yes |
| SM15.K05 | F | wild | -1.304 | 36.693 | 0 | 0 | PCR | yes |
| SM15.K06 | M | wild | -1.304 | 36.693 | 0 | 0 | PCR | no |
| SM15.K07 | F | wild | -1.304 | 36.693 | 0 | 0 | PCR | no |
| SM15.K08 | M | wild | -1.304 | 36.693 | 0 | 0 | PCR | no |
| SM15.K09 | M | wild | -1.304 | 36.693 | 0 | 0 | PCR | no |
| SM15.K10 | M | wild | -1.304 | 36.693 | 0 | 0 | PCR | no |
| SM15.K21 | F | wild | -1.304 | 36.693 | 1 | 1 | PCR | yes |
| SM16.L01 | M | wild | 0.302 | 36.908 | 0 | 0 | PCR | no |
| SM16.L02 | F | wild | 0.302 | 36.908 | 1 | 1 | PCR | yes |
| SM16.L03 | F | wild | 0.302 | 36.908 | 1 | 1 | PCR | yes |
| SM16.L04 | F | wild | 0.302 | 36.908 | 0 | 0 | PCR | no |
| SM16.L05 | F | wild | 0.302 | 36.908 | 1 | 1 | PCR | no |
| SM16.L06 | F | wild | 0.302 | 36.908 | 0 | 0 | PCR | no |
| SM16.L07 | F | wild | 0.302 | 36.908 | 1 | 1 | PCR | no |
| SM16.L08 | F | wild | 0.302 | 36.908 | 1 | 1 | PCR | no |
| SM16.L09 | M | wild | 0.302 | 36.908 | 0 | 0 | PCR | yes |
| SM16.L10 | M | wild | 0.302 | 36.908 | 0 | 0 | PCR | no |
| SM16.L11 | F | wild | 0.302 | 36.908 | 0 | 0 | PCR | yes |
| SM16.L12 | F | wild | 0.302 | 36.908 | 0 | 0 | PCR | no |
| SM16.L13 | F | wild | 0.302 | 36.908 | 0 | 0 | PCR | no |
| SM16.L14 | F | wild | 0.302 | 36.908 | 0 | 0 | PCR | no |
| SM16.L15 | M | wild | 0.302 | 36.908 | 0 | 0 | PCR | no |
| SM16.L16 | M | wild | 0.302 | 36.908 | 0 | 0 | PCR | no |
| SM16.L17 | F | wild | 0.302 | 36.908 | 0 | 0 | PCR | no |
| SM16.L18 | F | wild | 0.302 | 36.908 | 0 | 0 | PCR | no |
| SM16.L19 | F | wild | 0.302 | 36.908 | 0 | 0 | PCR | no |
| SM16.L20 | F | wild | 0.302 | 36.908 | 0 | 0 | PCR | no |

| | | | | | | | | |
|---|---|---|---|---|---|---|---|---|
| SM16.L21 | F | wild | 0.302 | 36.908 | 0 | 0 | PCR | no |
| SM16.L22 | M | wild | 0.302 | 36.908 | 0 | 0 | PCR | yes |
| SM16.L23 | M | wild | 0.302 | 36.908 | 0 | 0 | PCR | no |
| SM16.L24 | M | wild | 0.302 | 36.908 | 0 | 0 | PCR | no |
| SM16.L25 | M | wild | 0.302 | 36.908 | 0 | 0 | PCR | no |
| SM16.L26 | M | wild | 0.302 | 36.908 | 0 | 0 | PCR | no |
| SM16.L27 | M | wild | 0.302 | 36.908 | 0 | 0 | PCR | no |
| SM16.L28 | M | wild | 0.302 | 36.908 | 0 | 0 | PCR | no |
| SM16.L29 | F | wild | 0.302 | 36.908 | 0 | 0 | PCR | no |
| SM16.L30 | F | wild | 0.302 | 36.908 | 0 | 0 | PCR | no |
| SM16.L31 | M | wild | 0.302 | 36.908 | 0 | 0 | PCR | no |
| SM16.L32 | F | wild | 0.302 | 36.908 | 0 | 0 | PCR | no |
| SM16.L33 | F | wild | 0.302 | 36.908 | 0 | 0 | PCR | no |
| SM16.L34 | F | wild | 0.302 | 36.908 | 0 | 0 | PCR | yes |
| SM16.L35 | F | wild | 0.302 | 36.908 | 0 | 0 | PCR | no |
| SM16.L36 | M | wild | 0.302 | 36.908 | 0 | 0 | PCR | no |
| SM16.L37 | F | wild | 0.302 | 36.908 | 0 | 0 | PCR | no |
| SM16.L38 | F | wild | 0.302 | 36.908 | 0 | 0 | PCR | no |
| SM16.L39 | F | wild | 0.302 | 36.908 | 0 | 0 | PCR | no |
| SM16.L40 | M | wild | 0.302 | 36.908 | 0 | 0 | PCR | no |
| SM16.L41 | M | wild | 0.302 | 36.908 | 0 | 0 | PCR | no |
| SM16.L42 | F | wild | 0.302 | 36.908 | 0 | 0 | PCR | no |
| SM16.M03 | F | wild | 0.302 | 36.908 | 0 | 0 | PCR | no |
| SM16.N01 | F | wild | 6.24 | 8.97 | 0 | 0 | WGS | yes |
| SM16.N04 | F | wild | 6.24 | 8.97 | 0 | 0 | WGS & PCR | yes |
| SM16.N05 | M | wild | 6.24 | 8.97 | 0 | 0 | WGS | yes |
| SM16.N06 | M | wild | 6.24 | 8.97 | 0 | 0 | WGS | yes |
| SM16.N20 | M | wild | 6.24 | 8.97 | 0 | 0 | WGS | yes |
| SM16.N37 | M | wild | 6.24 | 8.97 | 0 | 0 | WGS | yes |
| DS98.324 | F | wild | 17.12 | 54.1 | 0 | 0 | PCR | no |
| SM17.F01 | F | wild | 28.331 | -13.923 | 0 | 0 | PCR | no |
| SM17.F02 | F | wild | 28.331 | -13.923 | 0 | 0 | PCR | no |
| SM17.F03 | F | wild | 28.331 | -13.923 | 0 | 0 | PCR | no |
| SM17.F04 | F | wild | 28.331 | -13.923 | 0 | 0 | PCR | no |
| RV12.N314 | U | wild | 35.549 | 9.754 | 0 | 0 | PCR | yes |
| RV12.N315 | U | wild | 35.549 | 9.754 | 0 | 0 | PCR | no |
| RV12.N317 | F | wild | 35.549 | 9.754 | 0 | 0 | WGS | yes |
| 18MX101 | F | reared | NA | NA | 1 | 1 | PCR | yes |
| 18MX102 | F | reared | NA | NA | 1 | 1 | PCR | no |

| | | | | | | | | |
|---|---|---|---|---|---|---|---|---|
| FJ97.1011 | F | reared | NA | NA | 1 | 1 | PCR | no |
| FJ97.1017 | F | reared | NA | NA | 1 | 1 | PCR | no |
| FJ97.1026 | F | reared | NA | NA | 1 | 1 | PCR | no |
| FJ97.1041 | F | reared | NA | NA | 1 | 1 | PCR | yes |
| FJ97.1283 | F | reared | NA | NA | 0 | 0 | PCR | yes |
| FJ97.1341 | F | reared | NA | NA | 0 | 0 | PCR | no |
| FJ97.466 | F | reared | NA | NA | 1 | 1 | PCR | no |
| FJ97.470 | F | reared | NA | NA | 1 | 1 | PCR | no |
| FJ97.491 | F | reared | NA | NA | 1 | 1 | PCR | yes |
| FJ97.633 | F | reared | NA | NA | 1 | 1 | PCR | no |
| SM17.P01 | F | reared | NA | NA | 0 | 0 | WGS | yes |
| SM17.X01 | F | reared | NA | NA | 1 | 0 | WGS | yes |
